# The regulatory landscape of cells in the developing mouse cerebellum

**DOI:** 10.1101/2021.01.29.428632

**Authors:** Ioannis Sarropoulos, Mari Sepp, Robert Frömel, Kevin Leiss, Nils Trost, Evgeny Leushkin, Konstantin Okonechnikov, Piyush Joshi, Lena M. Kutscher, Margarida Cardoso-Moreira, Stefan M. Pfister, Henrik Kaessmann

**Author notes:** These authors contributed equally to this study (Ioannis Sarropoulos, Mari Sepp). These authors jointly supervised the work (Stefan M. Pfister, Henrik Kaessmann). Correspondence (H.K.), (S.M.P.), (I.S.), (M.S.).

## Abstract

Organ development is orchestrated by cell- and time-specific gene regulatory networks. Here we investigated the regulatory basis of mouse cerebellum development from early neurogenesis to adulthood. By acquiring snATAC-seq profiles for ~90,000 cells spanning eleven stages, we mapped all major cerebellar cell types and identified candidate *cis*-regulatory elements (CREs). We detected extensive spatiotemporal heterogeneity among progenitor cells and characterized the regulatory programs underlying the differentiation of cerebellar neurons. Although CRE activity is predominantly cell type- and time-specific, periods of greater regulatory change are shared across cell types. There is a universal decrease in CRE conservation and pleiotropy during development and differentiation, but the degree of evolutionary constraint differs between cerebellar cell types. Our work delineates the developmental and evolutionary dynamics of gene regulation in cerebellar cells and provides general insights into mammalian organ development.

## Main text

The cerebellum is primarily known for its role in motor control, but it also contributes to other complex functions, such as language and memory (*1*–*4*). Abnormal development and malfunctioning of the cerebellum are linked to a series of human disorders, including ataxia, schizophrenia, autism, and pediatric brain tumors, such as medulloblastoma and ependymoma (*1*, *2*). The basic neural circuit of the cerebellum centers on the inhibitory Purkinje cells (PCs), which integrate information from other areas of the central nervous system and transmit signals to the neurons of the cerebellar deep nuclei (DN). The incoming information is modulated by a dense layer of excitatory granule cells (GCs) and a diverse set of inhibitory GABAergic interneurons (INs). Although this simple circuit is conserved across jawed vertebrate species, the size, morphology and developmental adaptations of the cerebellum show great variation (*4*, *5*). Such variation is especially pronounced in the mechanisms underlying the amplification and migration of GCs (*4*, *6*, *7*), which are the most abundant neuron type in the cerebellum, and – in mammals – in the entire brain (*8*).

Cerebellum development in mammals relies on a spatially and temporally restricted pattern of cell type specification. Two different germinal zones, the ventricular zone and the rhombic lip give rise to distinct neuronal and glial populations in a temporally restricted manner. GABAergic deep nuclei neurons (GABA DNs), PCs, INs and astrocytes are sequentially derived from progenitors in the ventricular zone, whereas the emergence of glutamatergic deep nuclei neurons (Glut. DNs) from progenitors in the rhombic lip is followed by the generation of GC progenitors (GCPs) and unipolar brush cells (UBCs) (*2*). Decades of molecular research have identified major regulators of these processes, including the transcription factors (TFs) ATOH1 and PTF1A, which are essential for the formation of the rhombic lip and ventricular zone, respectively (*2*, *9*). Two other TFs, OLIG2 and GSX1, are responsible for the temporal switch of the ventricular zone from PCs towards INs (*2*, *10*). Single-cell transcriptomics studies have provided additional cellular and molecular insights into cerebellum development (*11*–*15*) but the regulatory basis of these dynamic expression programs remains largely unexplored.

Developmental gene expression is largely controlled by the interplay between *cis*-regulatory elements (CREs), such as enhancers, silencers and promoters, and the TFs that bind to them (*16*, *17*). By integrating information from the binding of lineage-determining and signal-dependent TFs, the 3D chromosomal structure and the local chromatin state, distal CREs drive cell type-specific gene expression (*16*, *18*), and concomitant specification of cell type identity (*19*, *20*), thereby contributing to the precise control of organ development (*16*, *21*, *22*). Bulk measurements of CRE activity from the hindbrain (*22*, *23*), postnatal cerebellum and *in vitro* differentiating GCs (*24*) have provided important insights but lack the cellular resolution required to characterize the regulatory programs associated with specification and differentiation of cell types in the developing cerebellum.

Most CREs undergo rapid evolutionary turnover (*25*–*28*), contrary to the gene expression programs they control (*29*, *30*). However, a small number of regulatory regions, primarily those associated with developmental genes, show remarkable sequence and activity conservation, even comparable to that of protein-coding exon sequences (*31*, *32*). Previous studies analyzing whole tissues (*21*) and *in vitro* cultured cell lines (*20*) reported higher conservation levels of CREs active at early stages of organ development, which also show the strongest selective constraints in gene expression (*30*, *33*). However, potential differences in the regulatory conservation between cell types, which might have arisen due to different selective pressures and evolutionary histories, and their contributions to the previously observed whole-organ patterns, have so far not been systematically explored.

Recent methods based on the Assay for Transposase Accessible Chromatin (ATAC-seq) (*34*) enable the measurement of chromatin accessibility, a proxy for regulatory element activity, at the single-cell level (*35*– *37*). These have been employed to study gene regulatory activities in adult mouse organs (*38*), including the brain (*39*), as well as a limited number of cells or stages from the developing mouse (*40*, *41*) and human brain (*42*, *43*). In this study, we used single-nucleus ATAC-seq (snATAC-seq) to profile the chromatin accessibility landscape of ~90,000 cells across the development of the mouse cerebellum (https://apps.kaessmannlab.org/mouse_cereb_atac/). These data allowed us to identify cell type- and stage-specific patterns of CRE and TF activities and to characterize their contributions towards cell fate specification and differentiation in the developing cerebellum. We further investigated the evolutionary dynamics of CREs across cell types and stages, unveiling common trends, but also differences between cell types. Our work sheds light on the dynamics of gene regulation in the mouse cerebellum at the level of individual cell types, and provides further support for longstanding principles underlying the interplay between organ development and evolution.

## Results

### A single-cell chromatin accessibility atlas of the developing mouse cerebellum

We dissected mouse cerebella from 6 prenatal and 5 postnatal developmental stages, ranging from early neurogenesis on embryonic day 10 (E10) to adulthood (postnatal day 63; P63; Methods; Fig. 1A). We included two biological replicates per developmental stage, one from each sex (Table S1). We extracted nuclei from the entire cerebellum to ensure unbiased coverage of cell types, prepared snATAC-seq libraries using 10x Chromium (*36*), and then sequenced each library to an average depth of ~220 million read pairs. After applying strict quality control metrics (Methods) (*44*), including the *in silico* identification and removal of putative doublets (Fig. S1A-D), we acquired single-cell chromatin accessibility profiles for a total of 91,922 high quality cells with a median of 20,558 fragments per cell (Fig. 1B; Table S2). Profiles of biological replicates show the highest correlations (Spearman’s *rho* = 0.94-0.98), followed by correlations between samples from adjacent developmental stages (Fig. S1E). Using an iterative clustering approach and approximating gene expression based on the aggregated accessibility across a gene’s regulatory landscape (gene activity; Methods) (*44*), we identified 15 major cell types (including 9 neuronal cell types), which were further subdivided into 43 subtypes and cell states (Fig. 1C-E, S1F-H, S2A-B; Table S2).

**Fig. 1.**
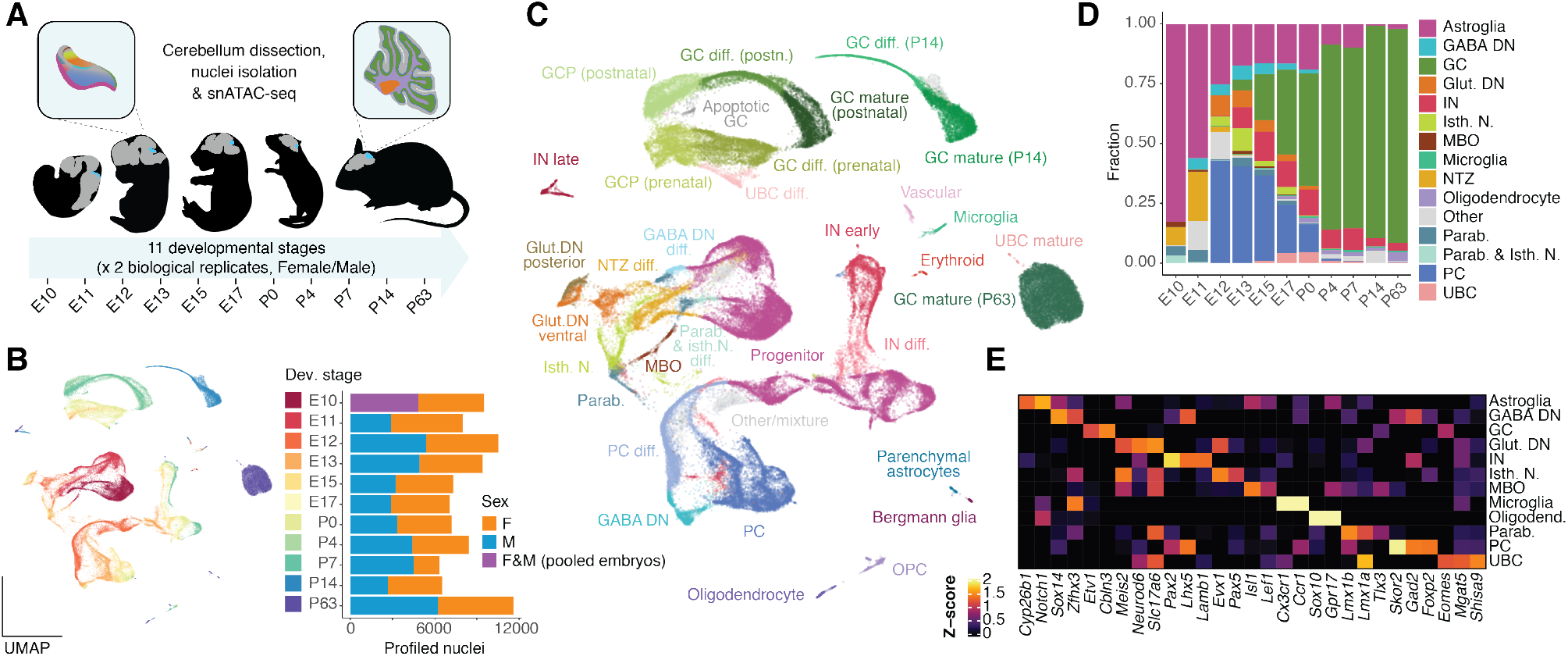
A chromatin accessibility atlas of mouse cerebellum development at single-cell resolution. (A) Schematic overview of the dataset. Representative mouse silhouettes are shown for E11, E13, E17, P4 and P63 (brain in grey, cerebellum in cyan). The insets show major cell types and their location in the cerebellum for E13 and P63. Cell type colors are as in (D). (B, C) UMAP projection of 91,922 high quality cells colored by developmental stage (B) or cell type and state (C). Barplots in B show the number of profiled cells per stage and sex (each sex corresponding to one biological replicate). Cell type abbreviations as in text; diff., differentiating; OPC, oligodendrocyte progenitor cell. (D) Proportions of broad cell types across developmental stages. Astroglia lineage includes neural progenitors and astrocytes. (E) Activity scores of genes used for the annotation of major cell types (Z-score, capped to 0-2). Detailed view of marker genes across subtypes and states in Fig. S2A.

We validated the use of gene activity as a proxy for gene expression and the quality of our cell type annotation through the stage-wise integration of our snATAC-seq data with a scRNA-seq atlas of the developing mouse hindbrain and cerebellum (*12*). Despite differences in dissections and sampled developmental stages, we observed a high concordance between the cell type labels assigned by the two studies (Fig. S2C), and good correlations between gene activities and integration-imputed RNA expression values (Fig. S2D). Minor discrepancies between the two annotations can be explained by some scRNA-seq clusters containing mixtures of multiple cell types (Methods; Fig. S2E-F).

The earliest developmental stages in our dataset (E10-E11) are dominated by neural progenitors (key marker genes: *Notch1, Cyp26b1*; Fig. 1C-E). We also detected midbrain-originating cells (MBO; *Isl1*) and parabrachial neurons (Parab.; *Lmx1b*), which migrate out of the cerebellum at later stages (*13*, *45*, *46*), as well as differentiating GABA DNs (*Zfhx3, Sox14*) and nuclear transitory zone (NTZ) glutamatergic neurons (*Meis2, Neurod6*). From E12 onwards, the NTZ could be further resolved into distinct posterior (*Lmx1a*) and ventral (*Lmo3*) populations of Glut. DNs, and into anteriorly located isthmic nuclei neurons (Isth. N.; *Pax5*) (*13*). The same developmental window is marked by the generation of a large number of PCs (*Skor2, Foxp2*), which are the most abundant neuron type until E15, gradually being outnumbered by INs (*Pax2*) and GCPs (*Atoh1*). During these first five days of cerebellum development, the relative abundance of progenitor cells (not including GCPs), decreases from 83% in E10 to 17% in E15 (Fig. 1D).

The next stages (E17-P4) are marked by the continuous generation of INs, the emergence of UBCs (*Lmx1a, Eomes*), and the rapid expansion of the proliferating (*Atoh1, Gli2*) and differentiating (*Neurod1, Grin2b*) GC populations, which account for 77% of the cerebellar cells in P4 (Fig. 1C-E). Small numbers of microglia (*Ccr1, Hexb*) and oligodendrocytes (*Sox10*), most of which are of extracerebellar origin (*2*), are also detectable from E17 onwards. The last two developmental stages (P14, P63) are dominated by mature GCs (*Etv1, Cbln3*; 89% of all cells in both stages), with additional neuronal and glial populations being traceable, including mature astroglia such as Bergmann glia (*Gdf10*) and parenchymal astrocytes (*Aqp4, Scl6a11*). Throughout this study, we use the term astroglia more generally to refer to cells transitioning along the lineage of neuroepithelial progenitors, radial glial progenitors and mature astrocytes (Fig. 1D-E), in agreement with their overlapping functions and molecular features (*47*, *48*). Overall, the developmental dynamics of cerebellar cell abundances observed in this study closely resemble those from single-cell RNA sequencing (scRNA-seq) atlases (*11*, *12*). However, some cell types, especially the early-born neurons (GABA DNs, glut. DNs, isth.N and parab.) are better resolved in our snATAC-seq atlas.

### The cis-regulatory landscape of cerebellar cell types

Having established a cellular atlas of mouse cerebellum development, we next sought to characterize the regulatory profiles of individual cerebellar cell types. We employed a cluster-specific and replicate-aware peak calling approach (*44*), identifying a total of 261,643 high-confidence putative CREs (Fig. S3A-B; Methods). 36,461 (14%) of these peaks overlap a protein-coding gene promoter or exon, with an additional 24,228 (9%) peaks being associated with a long non-coding RNA (lncRNA), small RNA, or other annotated transcript (Fig. 2A). However, the majority of the peaks are intergenic (67,630; 26%) or intronic (133,323; 51%) (herein collectively referred to as distal). Benchmarking this putative CRE (hereafter: CRE) set against external datasets (*38*, *49*, *50*) revealed a strong enrichment for putative enhancer activity during hindbrain development and high activity in cells from the adult cerebellum when compared with other organs (Fig. S3C-F).

**Fig. 2.**
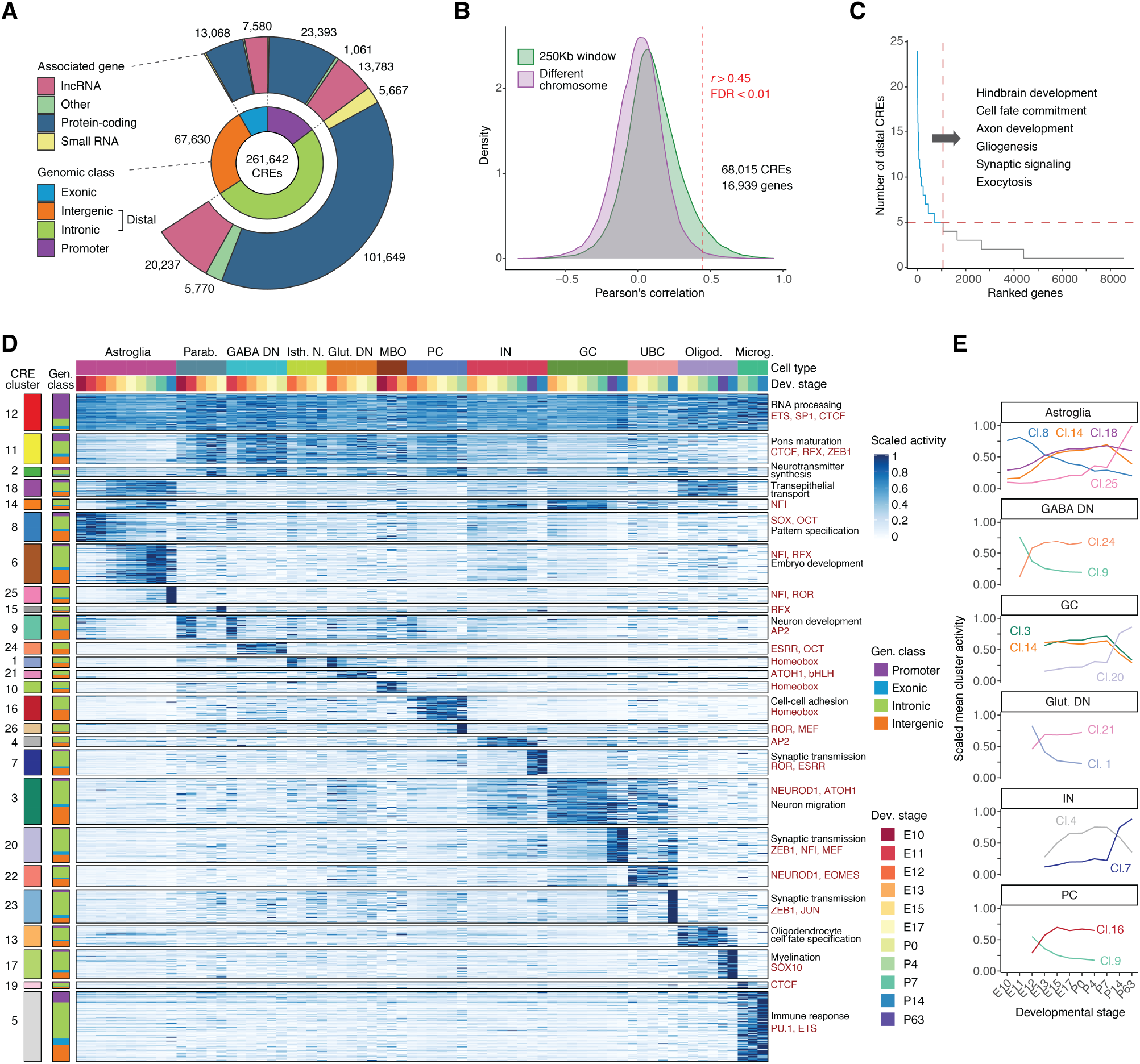
CRE identification, annotation and activity in cerebellum development. (A) Genomic features of 261,642 putative CREs. Inner and outer circles illustrate the genomic class of the CREs (inner) and the biotype of the associated genes (outer). (B) Density plot of Pearson’s *r* correlation coefficients between CRE accessibility and gene activity in +/-250 Kb windows considered for target assignment (green) and in different chromosomes used as control (purple). The red line indicates the cutoff used for assignment at a false discovery rate (FDR) < 1%. (C) Number of distal CREs per gene in decreasing order. Enriched biological processes for genes with five or more distal CREs are shown (BH adjusted *P* < 0.05; hypergeometric test). (D) Heatmap illustrating CRE activity (in counts per million, CPM, scaled by its maximum value) across cell types and developmental stages (top). CREs are grouped by activity cluster and genomic class (left). CRE clusters are arranged in decreasing order of pleiotropy (here: mean activity across rows) and then by cell type and developmental stage with maximum activity. Representative enrichments for biological processes of adjacent genes (black) and motifs for TFs or TF families (red) are shown on the right. 50,000 CREs confidently assigned to their cluster (Pearson’s *r* with cluster mean greater than 0.5) were chosen randomly for visualization purposes. (E) Mean scaled activity of CRE clusters active in different cell types across developmental stages.

We assigned CREs to their putative target genes based on correlations between gene activity and adjacent peak accessibility (Methods; Fig. 2B, S3A). More than 25% (68,015) of the CREs were assigned to at least one gene, with 80% of protein-coding genes showing dynamic expression during cerebellum development (*30*) being associated with at least one CRE (57% with a distal CRE). On average, each gene is associated with a single distal CRE, but 1,058 protein-coding genes were assigned to five or more distal CREs (Fig. 2C). These genes are enriched for developmental processes, such as cell fate commitment and neuron differentiation, as well as mature neuron functions like synapse organization and signaling (Fig. S3G).

We then used an iterative clustering procedure to identify the major patterns of CRE activity during cerebellum development (Methods). We identified 26 activity clusters, most of which are cell type- and time-specific (Fig. 2D-E). Cell type-specific CREs are close to genes associated with relevant gene ontology terms (e.g., myelination for oligodendrocyte-specific CREs) and enriched for motifs of TFs known to be active in the respective cell types (e.g., ATOH1 for GCs, SOX family TFs for progenitors, PU.1 for microglia; Fig. S3J-K). Although most CREs are cell type- and time-specific, we also identified a cluster showing constitutive activity (cluster 12; Fig. 2D) that is enriched for promoters as well as distal elements with CTCF motifs (Fig. S3H-I). We further observed groups of CREs active in multiple early-born neuron types (clusters 2, 11), glial populations (cluster 18) and late-born cell types (cluster 14), supporting the notion that a sizeable fraction of CREs shows pleiotropic activity (*51*, *52*).

### Periods of greater regulatory change

In our global clustering analysis (Fig. 2D), most CRE clusters are cell type-specific and characterized by sharp inflexion points in their activity during development (e.g., Fig. 2E). This suggests that specific developmental stages are associated with coordinated changes in the activity of multiple CREs within each cell type. To assess this hypothesis, we identified CREs with significant changes in accessibility between adjacent stages for the most abundant cerebellar cell types (Methods; Fig. 3A). Each cell type is characterized by a distinct set of differentially accessible CREs (Fig. 3B, S4A) and developmental pattern. For example, major changes in PCs occur one day after major changes in GABA DNs, reflecting the temporal order in which the two neuronal types are generated (Fig. 1D, 3A). Despite these differences, cell type-specific changes in CRE activity tend to be concentrated around the same developmental windows, leading to specific periods of greater regulatory change that can be detected at the level of the entire organ (Fig. 3C). Since our power to detect differentially accessible CREs depends on cell abundance, and thus library size (Methods; Fig. S4B), we repeated this analysis using the same number of fragments for each sample. Although we detected fewer differentially accessible CREs, we identified the same developmental windows to be associated with greater change (Fig. S4C). These periods were also marked by lower correlations in chromatin accessibility profiles of cell types across adjacent stages (Methods; Fig. S4D).

**Fig. 3.**
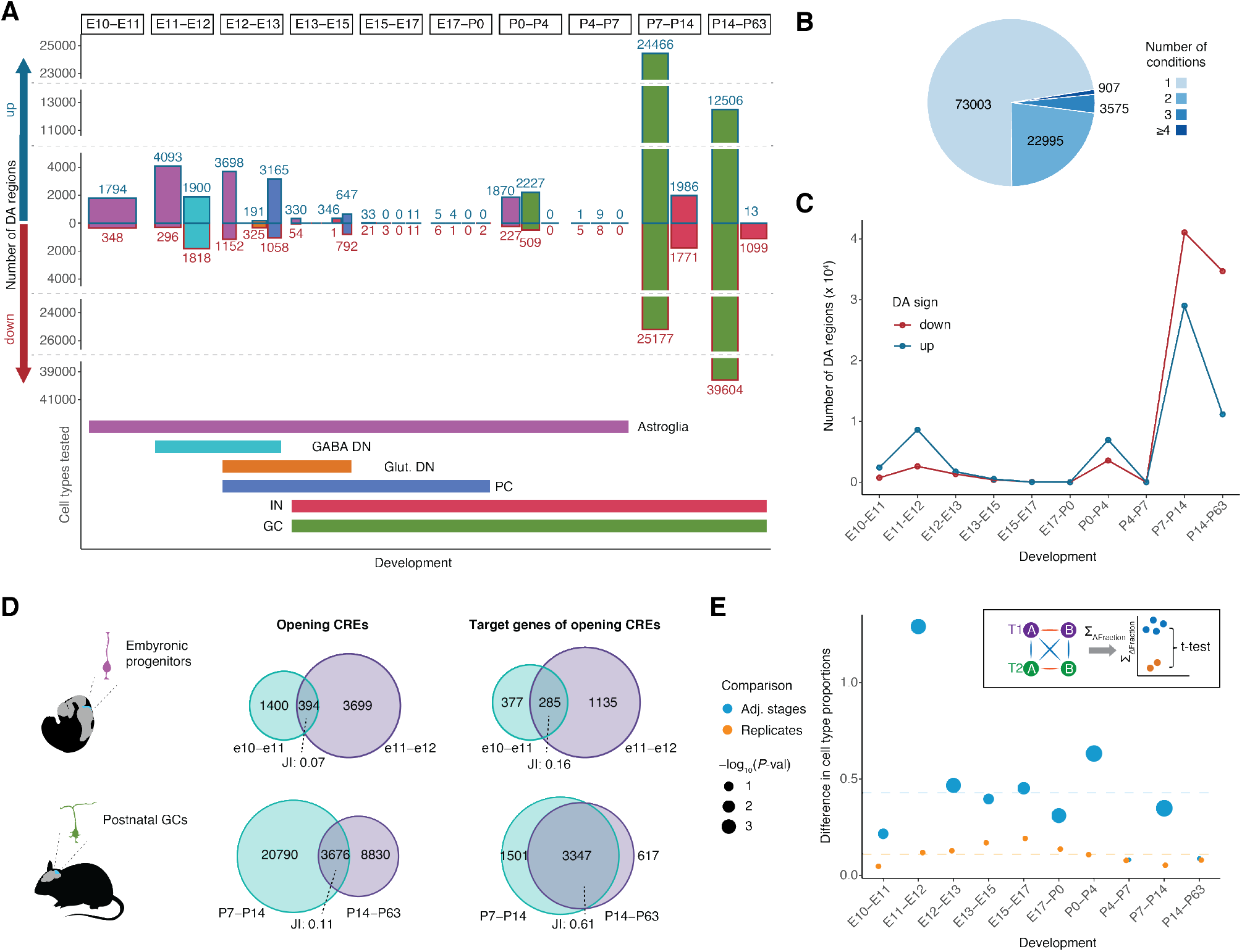
Periods of greater regulatory change. (A) Number of differentially accessible (DA) CREs across adjacent developmental stages in major cerebellar cell types (up: increasing accessibility; down: decreasing accessibility). Horizontal bars in the bottom indicate the developmental window for which each cell type was tested (restricted by its abundance). The y-axis is interrupted for visualization purposes. (B) Number of conditions (cell type and developmental stage) in which each CRE was identified as differentially accessible. (C) Number of differentially accessible CREs per developmental stage after aggregating the accessibility profiles across cell types. (D) Overlap between CREs (left) and putative target genes (right) with increased accessibility between adjacent stages in early progenitors (top) and postnatal GCs (bottom). Similarity across sets is quantified with Jaccard similarity indexes (JI). (E) Sum of absolute differences in cell type proportions across adjacent developmental stages (blue) and between biological replicates (orange). The size of the blue dots corresponds to the –log_10_ *P*-value of a t-test comparing differences across stages and between replicates. Horizontal lines correspond to mean values across stages. A summary of the method is illustrated in the inset. T1 and T2 correspond to consecutive stages, A and B to biological replicates, lines indicate pairwise differences in cell type proportions. Differences between (blue) and within (orange) stages are compared with a t-test.

During early embryogenesis (E10-E13), both progenitor cells (included in the astroglia lineage) and early-born neurons (GABA DNs, Glut. DNs and PCs) show major changes in their regulatory landscapes (Fig. 3A). In progenitor cells, CREs enriched for nuclear receptor motifs (NR2C2, NR2F2) are gradually replaced by regions enriched for POU and homeobox motifs (E10-E11), and eventually NFI motifs (E11-E12 and E12-E13; Fig. S4E). CREs proximal to genes involved in neuron differentiation and migration, and enriched for RFX binding sites, are activated in GABA DNs (E10-E11) and PCs (E11-E12; Fig. S4E-F). Despite this similarity, opening CREs in PCs also contain binding sites for homeobox TFs and RAR-related orphan receptors (RORs; Fig. S4E). This highly dynamic period of early organogenesis is followed by a period of fewer changes (E13-P0).

Immediately after birth (P0-P4), there is another period of greater regulatory change in the astroglia and GC populations, primarily involving the activation of CREs enriched for motifs of the NFI TF family in both cell types (Fig. 3A, S4E). Finally, major changes occur in GCs and INs between P7-P14 and P14-P63, leading to the closing of CREs near developmental genes and the opening of CREs adjacent to genes associated with ion transport and synaptic signaling (Fig. S4F). Unlike early progenitors (E10-E12), the putative target genes of CREs activated in postnatal GCs are largely shared between stages (Fig. 3D, S4F). However, despite this convergence in gene targets, the underlying differentially accessible CREs are distinct for each stage (Fig. 3B, 3D, S4A). CREs with increasing accessibility in P7-P14 are enriched for motifs of the ZEB, MEF and ROR TF families, whereas those activated in P14-P63 contain motifs recognized by basic leucine zipper domain (bZIP) factors, such as JUN (Fig. S4E). Thus, the activation of the same gene sets in postnatal GCs is regulated by TFs and CREs with temporally distinct activities. Furthermore, the fraction of intergenic elements and average distance to the closest transcription start site (TSS) decrease with time during these late postnatal stages (Fig. S4G-H), as previously observed for the developing forebrain (*21*).

Notably, the first two periods of greater regulatory change coincide with major alterations in the cellular composition of the cerebellum (emergence of PCs in E11-E12 and expansion of GCs in P0-P4; Fig. 3E, 1D). By contrast, few changes in cell type composition occur during later stages (P14-P63) of cerebellum development, which are dominated by GCs. Collectively, our analyses identified four highly dynamic periods during cerebellum development (E10-E13, P0-P4, P7-P14, P14-P63), characterized by greater changes in CRE activity across multiple cell types and, for the first two periods, in cellular composition.

### Heterogeneity of cerebellar progenitors reflects cell fate decisions

The observed temporal differences in cell type proportions (Fig. 1D, 3E) are largely due to the generation of distinct cell types in a spatially and temporally restricted manner (*2*, *53*). We asked whether this mode of cell fate specification was associated with spatiotemporal heterogeneity amongst cerebellar progenitors. Chromatin accessibility data are especially suitable for investigating this question, as regulatory changes often foreshadow gene expression during cell fate decisions (Ziffra et al., 2019; Ma et al., 2020) and are not obscured by cell cycle related genes, a major confounding factor in transcriptomics data (Ma et al., 2020). We thus subclustered cells from the astroglia lineage (progenitors and astrocytes). Our iterative clustering approach identified progenitors from all major germinal regions throughout cerebellum development, although without sharp boundaries within a given stage (Fig. 4A-B, S5A-C). Early progenitors (E10-E12) can be divided into isthmic (*En1, Pax5*), anterior ventricular zone (anterior VZ; *Gsx1, Wnt8b*), ventricular zone (VZ; *Dll1, Ptf1a* from E11) and rhombic lip (RL; *Cdon, Atoh1*) populations, as well as progenitors with no apparent commitment towards a cell fate (Fig. 4A-C, S5A-C), in accord with recent reports (*54*). E13 and E15 are marked by the appearance of two late progenitor populations (Fig. 4A-B) that broadly correspond to the previously described bipotent (*Gsx1, Wnt8b*) and gliogenic (*Slc1a3, Grm3*) progenitors (*55*). The bipotent progenitors migrate from the VZ to the prospective white matter (WM) and have been shown to generate interneurons and WM astrocytes; whereas the gliogenic progenitors locate to the developing Purkinje cell layer and give rise to Bergmann glia and granule cell layer (GCL) astrocytes (*55*, *56*). In line with this, we detected two populations of differentiating parenchymal astrocytes, astroblast WM (*Slc6a11*, *Olig2, Kcnd2*) and astroblast GCL (*Aqp4, Tekt5*) at late prenatal and early postnatal stages (E15-P7; Fig. 4A-C, S5A-C).

**Fig. 4.**
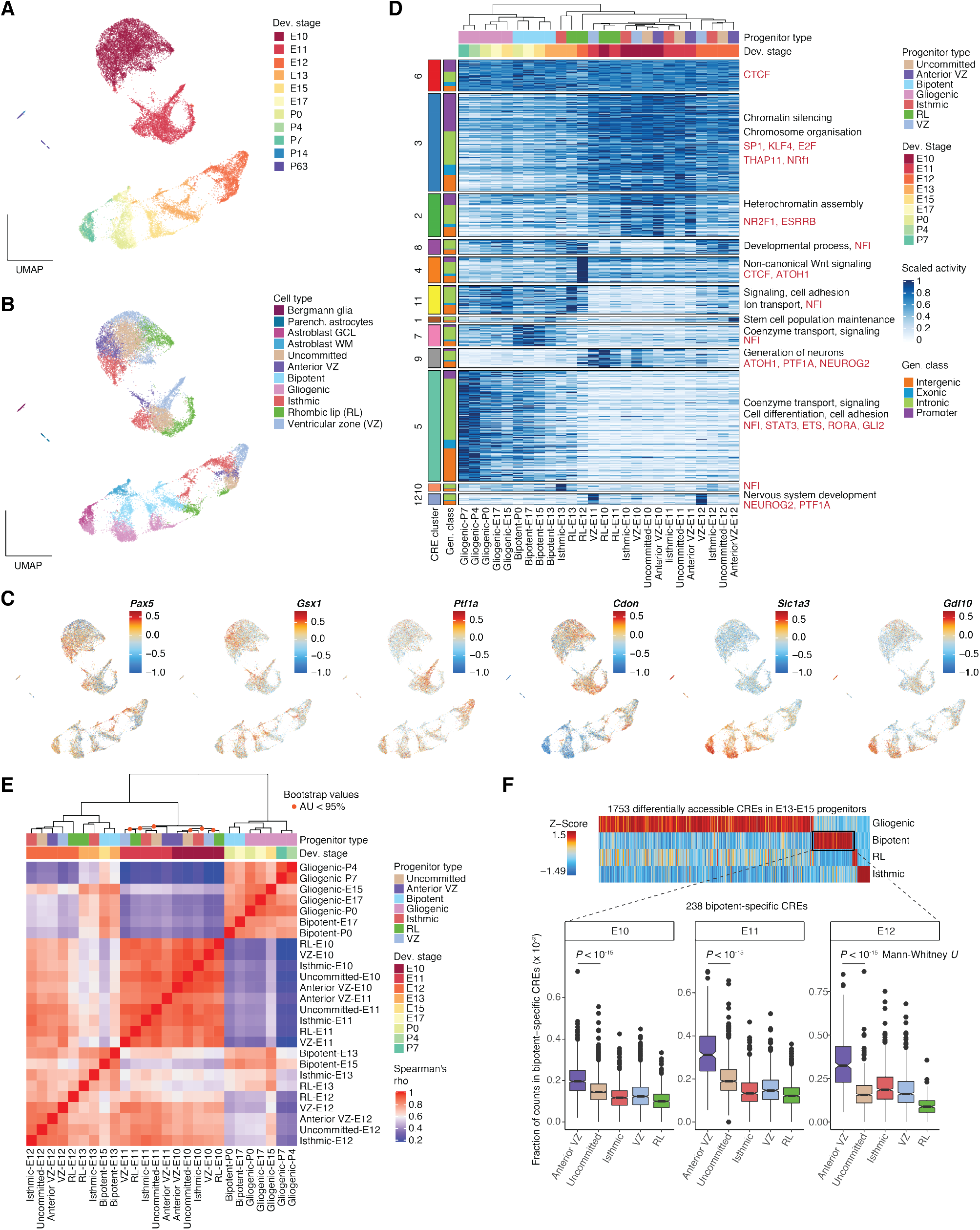
Spatiotemporal heterogeneity in cerebellar progenitor populations. (A, B) UMAP projections of 21,830 astroglia cells (progenitors and astrocytes) colored by their developmental stage (A) and cell type (B). (C) Gene activity scores for marker genes used for the annotation of progenitor populations (Counts per 10^4^ fragments, capped at 10^th^ and 99^th^ quantiles and log_10_ transformed). (D) Heatmap illustrating CRE activity (CPM scaled by its maximum value) across progenitor types and developmental stages (top). CREs are grouped by activity cluster and genomic class (left). Representative enrichments for biological processes of adjacent genes (black) and TF motifs (red) are shown on the right (BH adjusted *P* < 0.05; hypergeometric test). 50,000 CREs confidently assigned to their cluster (Pearson’s *r* with cluster mean greater than 0.5) were chosen randomly for visualization purposes. (E) Hierarchical clustering and heatmap of Spearman’s correlation coefficients in CRE accessibility across progenitor types and developmental stages. (F) Fraction of fragments in bipotent progenitor-specific CREs (E13-E15) for different progenitor populations and developmental stages.

Surprisingly, progenitor cells cluster primarily by developmental stage and not by germinal zone (Fig. 4A-B, E), despite our dense sampling of developmental stages and ability to identify the same germinal regions through consecutive stages, often on the basis of the same marker genes (e.g., RL in E10-E12; Fig. S5A). These temporal signals are strongest in early embryonic stages (E10-E12), a period coinciding with the generation of a diverse set of neuronal cell types (Fig. 4E, 1D). For example, within these two days, the VZ sequentially gives rise to parabrachial neurons, which exit the cerebellum towards ventral brainstem, GABA DNs, and PCs (*2*, *45*). Indeed, many CREs show temporally dynamic activity patterns that are shared between germinal zones (Fig. 4D), arguing for a mode of cell fate induction through shared temporal cues (*4*, *9*, *57*).

From E15 onwards, temporal differences become smaller and progenitors from adjacent stages group by germinal region (gliogenic and bipotent; Fig. 4E). Notably, this period coincides with the protracted generation of INs and astrocytes. Thus, although spatiotemporal heterogeneity among the progenitors is prominent throughout cerebellum development, early stages are characterized by stronger temporal signals, in agreement with the large number of differentially accessible CREs we identified in the same developmental window (Fig. 3A), whereas spatial signals increase towards late prenatal and postnatal stages.

A major temporal transition in cerebellar cell fate specification occurs in the VZ around E13 when *Olig2+* PC progenitors switch to *Gsx1+* bipotent progenitors that give rise to INs and eventually astrocytes (*10*). However, we identified a *Gsx1+* progenitor population already at E10, three days before we detected the first INs (Fig. 4A-C, S5A-C), consistent with an earlier study (*10*). Using *in situ* hybridization data (*58*) we confirmed the presence of these progenitors in the anterior VZ of the E11 cerebellum, between the PC-generating posterior VZ and the isthmus (Fig. S5B). Although this early *Gsx1+* progenitor population was previously observed (*10*), its relationship to the bipotent progenitors remains elusive. We assessed the similarity of these temporally distinct *Gsx1+* progenitor populations and found that additional marker genes, such as *Ndnf*, *Robo1* and *Wnt8b,* are shared between them (Fig. S5D). Furthermore, anterior VZ progenitors from E10-E12 showed the highest similarity in CRE activity to bipotent progenitors from E13 and E15, both globally and when focusing on cell type-specific CREs, across stage-matched progenitor types (Fig. 4F, S5E). Thus, our data provide support that early *Gsx1+* anterior VZ progenitors are developmentally related to the *Gsx1+* bipotent progenitors from later developmental stages, generating a hypothesis for future lineage tracing studies (*59*).

### Evolutionary dynamics of CREs across cell types and developmental stages

Motivated by the pronounced changes in CRE activity we observed during development, both in specified cell types (Fig. 3A) and within progenitor populations (Fig. 4A-E), we next investigated whether such temporal differences are also reflected in the evolutionary histories of CREs. A temporal decline in gene regulatory (*21*) and expression (*30*, *33*) conservation has been reported for whole organs, but differences between cell types, and their contributions to these developmental patterns, have not been explored. Hence, we decided to test for differences in the evolutionary dynamics of CREs both across cell types and throughout development.

We assessed the selective (functional) constraints on CRE sequences using estimates of evolutionary conservation (phastCons scores) (*60*), and inferred their minimum age (i.e., when they first appeared) using syntenic sequence alignments between high-quality genomes from mouse and 16 other vertebrates at various phylogenetic distances (Fig. S6A; Table S5; Methods). We then assessed these regulatory conservation metrics at the single-cell level (Methods; Fig. S6A-F), enabling comparisons between different cell types and developmental stages. Across all cell types, both sequence constraints and the predicted ages of distal (intronic and intergenic) CREs decreased significantly during development (Fig. 5A-B and S6G). This shows that the previous observations for whole organs (*21*) extend to individual cell types. Ancient CREs, shared across or even beyond mammals, are primarily active during embryonic development, when cell types are specified and begin their differentiation (Fig. 5E). As cell types mature and activate their terminal effector genes, elements specific to eutherians and murid rodents (Muridae) gradually increase their activity, potentially contributing to species/lineage-specific phenotypic innovations of ancestral cell types (Fig. 5C, E). Consistent with these observations, TF genes, which are central to cell type identity (*19*), are associated with older and more constrained CREs (Fig. 5D, *P* < 10^−15^, Mann-Whitney *U* test).

**Fig. 5.**
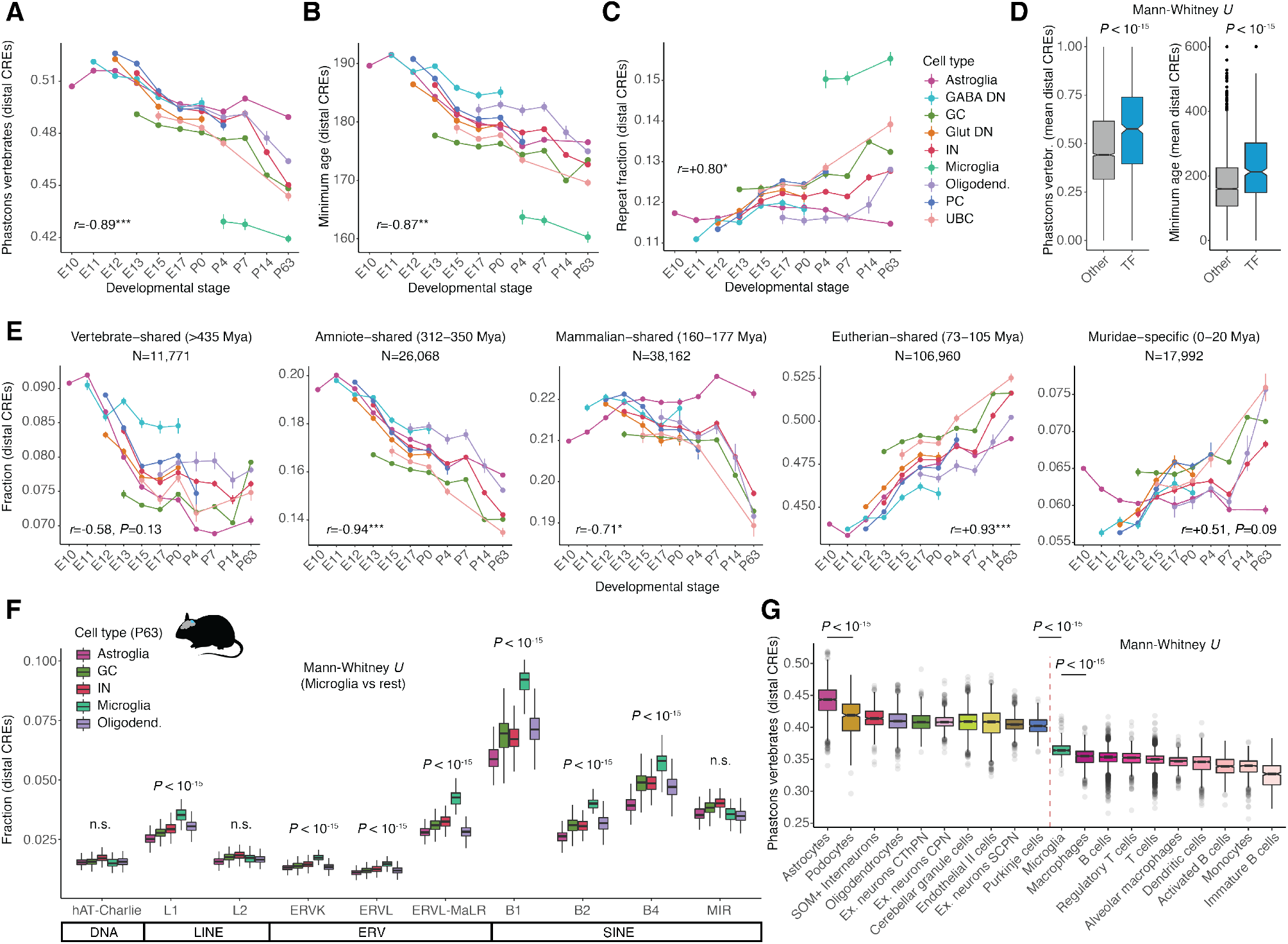
Evolutionary dynamics of CREs across cell types and developmental stages. (A, B, C) Average sequence constraint (A), inferred age (B) and fraction covered by repeats (C) in distal CREs active across cell types and developmental stages. Vertical bars illustrate 95% confidence intervals of the estimates. Pearson’s *r* correlation coefficients between the estimates and development are shown (median across cell types; *P*<0.05*, *P*<0.01**, *P*<0.001***). (D) Sequence constraint (left) and inferred age (right) of distal CREs associated with TFs and other genes with dynamic expression in mouse cerebellum development. (E) Fraction of fragments in distal CREs of different age groups per cell type and stage (CRE numbers indicated on top). Vertical bars illustrate 95% confidence intervals of the estimates. Pearson’s *r* correlation coefficients between the estimates and development are shown (median across cell types; *P*<0.05*, *P*<0.01**, *P*<0.001***). (F) Fraction of fragments in distal CREs overlapping TEs of different classes in cell types of the adult mouse cerebellum (P63). (G) Average sequence constraint of distal CREs active in different cell types of the adult mouse. The ten most conserved (left) and all immune (right) cell types are shown. Data, cell type and CRE annotations are from Cusanovich et al. 2018.

We also observed pronounced differences in the conservation of distal CREs between cell types. Most strikingly, CREs active in microglia show a much faster evolutionary turnover compared with all other cerebellar cell types (Fig. 5A-B and S6G; *P* < 10^−15^, Mann-Whitney *U* test), in line with the rapid divergence in gene expression and morphology of microglia (*61*). In agreement with their younger age and reduced sequence constraint, CREs active in microglia are also enriched for genomic repeats (Fig. 5C). This enrichment is driven by recently expanded transposable elements in rodents, such as short interspersed nuclear elements (SINEs) B1, B2 and B4, endogenous retrovirus sequences (ERVs), and L1 elements, and not by more ancient sequences shared across mammals (Fig. 5F). By contrast, astroglial cells (progenitors and astrocytes) show the most conserved distal CREs in the mature cerebellum (Fig. 5A and S6G; *P* < 10^−15^, Mann-Whitney *U* test) and constitute the only cell population without an increasing contribution of Muridae-specific and repeat-derived CREs during development (Fig. 5C, E). Instead, postnatal astroglia are enriched for distal CREs that originated 160-177 million years ago in common mammalian and therian (i.e., eutherian-marsupial) ancestors (Mya; Fig. 5E, *P* < 10^−15^, Mann-Whitney *U* test) and have since been strongly preserved by purifying selection across eutherian mammals (Fig. S6G), overall suggestive of a regulatory innovation in astroglia around the emergence of the major mammalian lineages. We confirmed and extended these observations using a single-cell chromatin accessibility atlas of adult mouse organs (*38*). Notably, we found that astrocytes have the most conserved distal CREs, not only in the adult cerebellum but also across all adult organs (Fig. 5G; *P* < 10^−15^, Mann-Whitney *U* test). Moreover, eight of the ten most conserved cell types across all organs were neural (Fig. 5G), highlighting the overall stronger evolutionary constraints in the brain (*30*, *62*). Consistently, despite having the most rapidly evolving regulatory landscape in the cerebellum, microglia constitute the most conserved immune cell type (Fig. 5G).

Among neurons, we also observed a decrease in the conservation of distal CREs during development, but the differences between cell types are subtler than among glial cells (Fig. 5A-B, S6G). In particular, GCs and UBCs exhibit significantly lower conservation levels compared with stage-matched Glut. DNs, GABA DNs, PCs and INs, especially during prenatal development (Fig. 5A, B; E13: *P* < 10^−15^, Mann-Whitney *U* test). This difference might reflect the overall high turnover in developmental adaptations associated with GC amplification across vertebrates (*4*), including the emergence of the external granule cell layer, a secondary germinal zone of proliferating GCs that is thought to be specific to amniotes (*6*, *7*). In agreement, developing GCs are depleted from ancient regulatory elements (shared across or beyond mammals) and enriched for eutherian and Muridae-specific elements (Fig. 5E). As in microglia, distal CREs in embryonic GCs (E13) are enriched for ERV and L1 transposable elements. However, in contrast to microglia, the most enriched SINE elements are mammalian interspersed repeats (MIRs), which were amplified before the mammalian radiation (*63*), with little to no enrichment for elements with rodent-specific expansion, such as B1 and B2 (Fig. S6H). Collectively, our results reveal common temporal trends as well as cell type-specific differences in the evolutionary dynamics of the *cis*-regulatory landscape of the developing cerebellum.

### Regulatory activity during neuron differentiation and maturation

We next sought to assess the contribution of gradual processes, such as cell type differentiation and maturation, to the observed temporal differences in CRE activity and conservation. We used diffusion pseudotime (*64*) to model these processes for the three major neuron types of the cerebellum: GCs, PCs and INs. We integrated data across multiple developmental stages starting immediately after cell fate commitment; that is, when cells could be unambiguously assigned to specific cell types (Methods, Fig. S7A-C). As expected, average pseudotime values increase with developmental stage (Fig. 6A-C). We observed a considerable overlap of differentiation states across stages for GCs and INs consistent with their protracted differentiation dynamics that span several weeks and expand into postnatal development (*2*). While GCs appear as a largely homogeneous differentiating population, INs are clearly stratified into distinct, temporally-specified subtypes (Fig. 6B, S7D-E). These include early-born INs (*Zfhx4, Slit2*) detected at E13-E15, mid-born Golgi (*Chrm2*), and Purkinje layer INs (PLI; *Nxph1, Klhl1*) prevalent at E17-P7, and late-born molecular layer INs of type 1 (MLI1; *Sorcs3, Grm8*) and 2 (MLI2; *Nxph1, Pvalb*), which are abundant at P14-P63 (Fig. S7D-E) (*2*, *65*). Contrary to the protracted trajectory of GCs and INs, PCs undergo rapid differentiation, primarily during E12 and E13, and then remain in a mostly steady differentiation state (Fig. 6C).

**Fig. 6.**
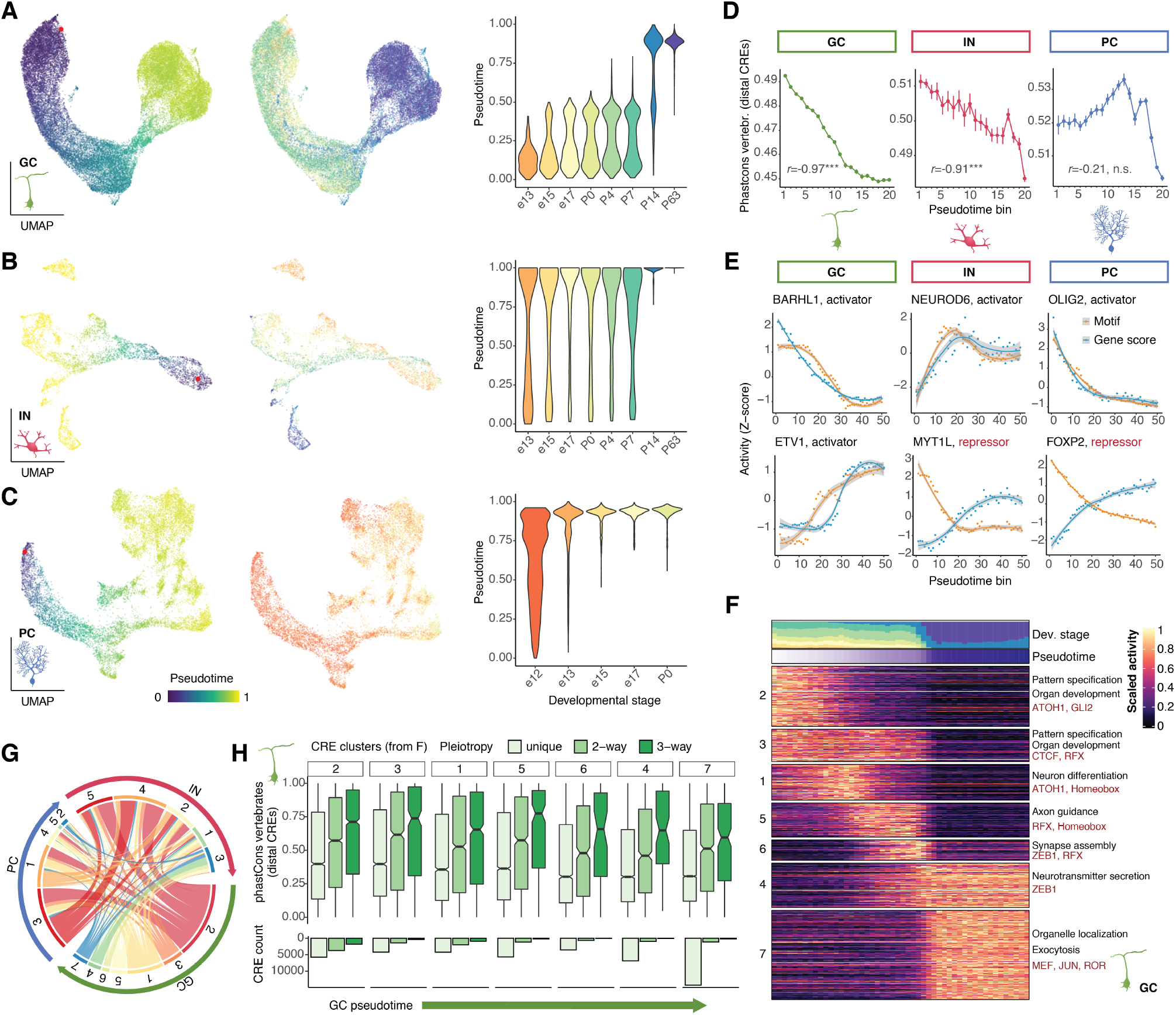
Regulatory landscape of neuronal differentiation in the developing cerebellum. (A, B, C) UMAP projections of 35,153 GCs (A), 5,113 INs (B) and 13,214 PCs (C) following integration across developmental stages. Cells are coded by pseudotime value (left) and developmental stage (middle). The violin plots (right) show the distribution of pseudotime values across developmental stages. The red points in the left panels indicate the pseudotime root. (D) Average sequence constraint of distal CREs across neuron differentiation. Cells are grouped in 20 bins in pseudotime intervals with a step of 0.05. Vertical bars illustrate 95% confidence intervals of the estimates. (E) Examples of TFs classified as activators and repressors. The points mark Z-score scaled values for gene (blue) and motif (orange) activity. Cells are grouped in 50 bins based on pseudotime ranks. Curves are drawn using LOESS regression and gray areas indicate 95% confidence intervals. (F) Heatmap of Z-score scaled activity of dynamic CREs across GC differentiation. Cells are grouped in 50 bins based on pseudotime ranks. The contribution of different developmental stages and mean pseudotime value for each bin are shown above the heatmap. Representative enrichments for biological processes of adjacent genes (black) and TF motifs (red) are shown on the right (BH adjusted *P* < 0.05; hypergeometric test). CRE clusters are shown on the left. (G) Chord diagram illustrating the overlap between activity clusters for CREs dynamic in two or more neurons (pleiotropic). The outer circle is colored by the neuron type. The inner circle indicates the activity clusters, indexed as in F and ordered from early (red, orange) to late (green, blue) activity. Each node represents a pairwise overlap between clusters from different neuron types. (H) Differences in sequence conservation (top) and abundance (bottom) of distal CREs with increasing pleiotropy (shading) active in different stages of GC differentiation (indicated by cluster numbers, as in F).

We then used the modeled trajectories to assess the conservation of distal CREs across differentiation. The sequence constraint decreases linearly during differentiation of GCs and INs, whereas in PCs it peaks at an intermediate state and then declines (Fig. 6D). Thus, differentiation signals modeled by pseudotime are mostly consistent, but not always identical, with temporal patterns observed across development (Fig. 5A).

Next, we characterized the gene regulatory dynamics along the differentiation trajectories of these three cerebellar neuron types. For each neuron type we identified TFs with dynamic activity during differentiation and integrated the accessibility of the respective binding motifs (*66*) with their inferred expression levels to classify TFs into putative activators and repressors (*67*) (Fig. S7F-H; Table S3; Methods). These candidate TFs include known regulators of neuronal differentiation, such as BARHL1 and ETV1 in GCs, MYT1L and NEUROD6 in INs, and OLIG2 and FOXP2 in PCs (Fig. 6E). At the level of individual CREs we observed a gradual transition from CREs proximal to genes associated with embryonic development and cell fate commitment, towards genes associated with neuron differentiation, followed by genes with roles in axon guidance and neuron migration across all three neuron types (Fig. 6F, S7I-J). CREs active at later pseudotime states are proximal to genes involved in the formation of the synapse and, eventually, in neurotransmitter secretion and ion transport (Fig. 6F, S7I-J). In support of this convergence in biological processes, 43% of protein-coding genes with dynamic activity across pseudotime are shared by at least two neuron types (Fig. S7K), in accordance with similar observations in the developing neocortex (*68*). By contrast, only 20% of dynamic CREs are shared across neuron types (i.e., they are pleiotropic), further highlighting the pronounced cell type-specificity of the *cis*-regulatory landscape (Fig. S7L). Pleiotropic CREs often show similar activity profiles across neuron types (Fig. 6G; early-early: *P* < 10^−15^, late-late: *P* < 10^−15^, early-late: *P* = 1, hypergeometric test) and are primarily active in the early stages of neuron differentiation (Fig. 6H, S7M-N). The higher similarity of the early stages is also supported by principal component analysis (PCA) in which the regulatory landscapes of the three neuron types gradually diverge as they differentiate (Fig. S7O). Pleiotropic CREs are also significantly more conserved than those that are dynamic in a single neuron type throughout all stages of differentiation (Fig. 6H, S7M-N). These results support the notion that pleiotropy imposes major constraints on the evolution of regulatory elements (*51*, *69*). As pleiotropic CRE activity decreases during development and differentiation, so does the degree of regulatory constraint, in agreement with previous observations for gene expression (*30*, *33*).

### Temporal signals in GC differentiation

Although we found that the dynamics of CRE activity and conservation across development can mostly be explained by cellular differentiation and maturation, additional temporal differences could be present, even between cells at the same differentiation state. Such differences could arise from intrinsic temporal patterning cues, as recently described in the context of cell fate specification (*57*, *68*), as well as extrinsic factors, such as changes in the availability of a morphogen or a ligand, synaptic simulation and interactions with neighboring cells (*16*). To assess the contribution of such temporal signals we focused on GCs, because these cells have a protracted differentiation trajectory (E13-P14) and do not have distinct temporally-specified subtypes. By stratifying GCs based on their differentiation state (pseudotime) and developmental stage, we observed that both factors contribute to the decrease of conservation of distal CREs during development, with cells at early differentiation states and developmental stages showing the most conserved regulatory program (Fig. 7A, S6I-J). To further characterize these differences, we divided the differentiating GCs into broad bins along the pseudotime (Fig. 7B) and identified differentially accessible CREs between cells from adjacent developmental stages within the same differentiation bin. In line with our previous observations (Fig. 1B, 3A), most radical changes could be detected postnatally, between P0-P4, P7-P14 and P14-P63 (Fig. 7C). These changes include opening of CREs near genes associated with neuron projection and synaptic ion transport in proliferating and migrating GCs (P0-P4), respectively, as well as downregulation of developmental genes in mature GCs (P14-P63; Table S4). In all differentiation state-matched comparisons, distal CREs closing during development are under stronger constraint than opening distal CREs (Fig. 7D).

**Fig. 7.**
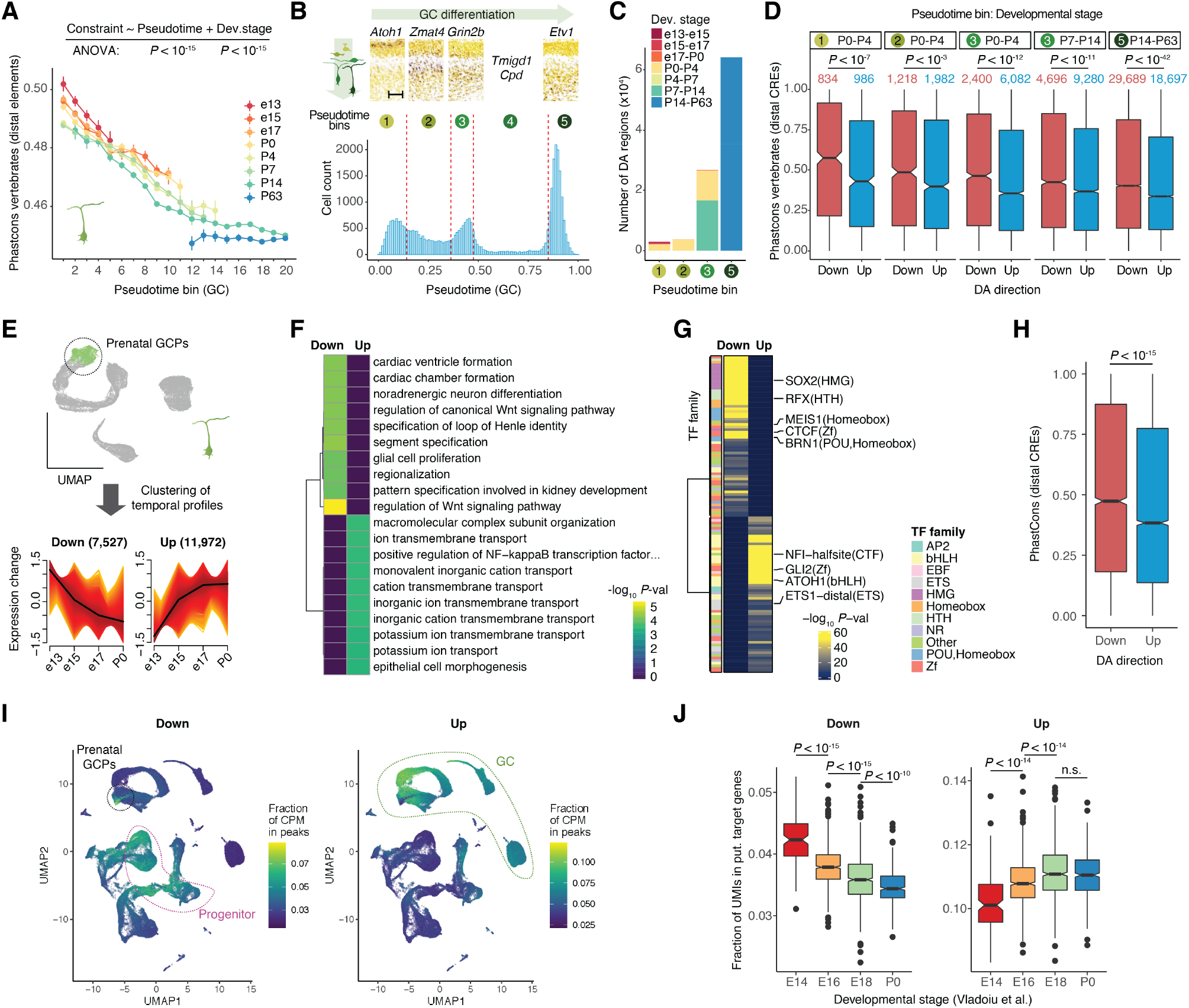
Temporal differences across matched differentiation states in GCs. (A) Average sequence constraint of distal CREs across GC differentiation. Cells are separated by developmental stage and grouped in 20 bins in pseudotime intervals with a step of 0.05. Vertical bars illustrate 95% confidence intervals of the estimates. (B) Distribution of pseudotime values across 35,153 GCs (left). Red vertical lines show the breaks for the five broad differentiation bins (also estimated based on cluster inflexion points). P4 *in situ* hybridization data of selected differentiation bin marker genes are shown above (*58*). Gradual shift of expression signals from external to internal granule layer of the cerebellar cortex is observed. Scale bar: 50 μm. (C) Number of differentially accessible (DA) CREs across adjacent developmental stages identified for each differentiation bin (as in B). (D) Sequence conservation of distal CREs opening (blue) and closing (red) across adjacent developmental stages only considering cells from the same differentiation bin (as in B). Numbers above the boxplots indicate the number of differentially accessible distal CREs. (E) Identification of temporal differences in CRE activity in prenatal GCPs (top; UMAP of 35,153 GCs prior to alignment across developmental stages, as in S7A). Z-score scaled temporal activity of opening and closing CREs (bottom). The black lines indicate the mean values of the clusters. (F, G, H) Characterization of developmentally dynamic CREs in prenatal GCPs in terms of enriched (BH adjusted P < 0.05; hypergeometric test) biological processes for adjacent genes (F), TF motifs (G) and sequence constraint (H). (F) and (G) show enrichments for all CREs, (H) shows constraint only for distal CREs. (I) Fraction of CPMs per cell in CREs that are closing (left) or opening (right) during development in prenatal GCPs. (J) Fraction of UMIs per cell in putative target genes of CREs that are closing (left) or opening (right) during development in prenatal GCPs. Data and annotations are from (*12*). *P*-values in (D), (H) and (J) are estimated by two-sided Mann-Whitney *U* tests.

We then focused on prenatal GCPs, which formed a single cluster prior to any alignment across developmental stages, and are thus at a similar differentiation state (Fig. 7E, S7A). This allowed us to control for potential artifacts caused by our inference of differentiation through the pseudotime analysis. Despite the overall high similarity of prenatal GCPs, we identified CREs with significant temporal activity changes (Methods) and classified them into two major clusters of decreasing (7,527) and increasing (11,972) accessibility across prenatal developmental stages (Fig. 7E). CREs with decreasing activity are more conserved than those with increasing activity and are associated with developmental functions and enriched for motifs recognized by pluripotency TFs, such as SOX2 and BRN1 (Fig. 7F-H). In contrast, CREs with increasing activity are associated with ion transport and enriched for motifs of TFs active in GCPs, such as ATOH1 and GLI2 (Fig. 7F-H). We also observed that closing CREs are active in other types of progenitor and early differentiating cells, whereas opening CREs show the highest activity in postnatal GCPs (Fig. 7I), further supporting the link between CRE pleiotropy and evolutionary constraint that we observed in the pseudotime analysis (Fig. 6H, S7M-N).

Since CREs with decreasing activity in the prenatal GCPs (E13-P0) are also active in progenitor cells from earlier developmental stages (E10-E12), we asked whether they still have an effect on gene expression or whether we were capturing the delayed compaction of already inactive CREs (*24*). To investigate this question, we considered the expression of their putative target genes in a scRNA-seq atlas of cerebellum development (*12*). We used proximity to assign CREs to genes to avoid the circularity introduced by our correlation-based approach, which requires genes and CREs to show similar patterns. The aggregate expression of genes adjacent to closing CREs decreased significantly in GCPs across prenatal developmental stages (Fig. 7J), suggesting that the progressive decline in CRE accessibility has an effect on gene expression. Collectively, our analyses provide evidence for temporal differences in CRE activity and gene expression in GCs that are not explained by differentiation and maturation processes.

## Discussion

In this study, we profiled ~90,000 cells to generate a regulatory atlas of the mouse cerebellum spanning eleven developmental stages, from the beginning of neurogenesis to adulthood. We showed that all major cerebellar cell types could be detected on the basis of their chromatin accessibility landscape. We identified candidate CREs for each cell type, most of which show cell type- and time-specific activity, and assigned them to putative target genes. We characterized the differentiation and maturation dynamics of the three major neuron types in the cerebellar circuit (PCs, GCs and INs), and identified CREs and TFs associated with these processes. Thus, we provide a comprehensive resource that will facilitate further research on the impact of gene regulation in cerebellum development and disease (https://apps.kaessmannlab.org/mouse_cereb_atac/).

### Temporal signals in CRE activity within cell types

Throughout our analyses we detected strong temporal differences in CRE activity within individual cell types. These are most apparent in the four periods of greater regulatory change (E10-E13, P0-P4, P7-P14, P14-P63). Although the first two periods are also associated with changes in the cellular composition of the cerebellum, the large number of radical changes identified at the cell type level suggests that the rapid temporal turnover in CRE activity previously described for whole organs (*21*) is mostly driven by changes within cell types. Despite being reminiscent of periods associated with major changes in the expression of coding and noncoding transcripts in the cerebellum and other mammalian organs (*30*, *33*), changes in regulatory activity seem more pervasive than at the level of the transcriptome. For example, very few radical expression changes occur in the cerebellum between P14 and P63 (*30*, *33*), while we observed numerous changes in CRE accessibility in this developmental window. The large number of CREs with decreasing accessibility in P14-P63 could be explained by the delayed closing of already deactivated developmental enhancers (*24*), whereas the high overlap between the putative target genes of opening CREs in P7-P14 and P14-P63 supports the idea that the expression of terminal effector genes in postmitotic neurons is maintained by the successive activation of transient enhancers (*70*).

Another manifestation of strong temporal signals was observed in cerebellar progenitors. Although we detected pronounced spatiotemporal heterogeneity across progenitor populations, we rarely identified CREs that were specific to a single germinal zone and developmental stage. Instead, our analyses suggest that this heterogeneity is driven by a combination of spatial and temporal signals. This in turn provides support for a model of cell fate induction, where different germinal zones jointly experience common temporal cues. These could involve intrinsic patterning through a shared temporal code, as recently suggested for multiple regions of the central nervous system (*57*). In support of this hypothesis, we found that late progenitor populations (E13-P7) are enriched for motifs of the NFI TF family, which is associated with the generation of late-born neurons in several brain regions (*57*). Additionally, common extrinsic cues, such as factors secreted from the choroid plexus might also explain the coordinated changes in CRE activity across germinal zones (*4*).

We also observed temporal signals in GCs that could not be explained by differentiation and maturation. Although these temporal differences in CRE activity have an effect on gene expression, their functional impact remains unclear. They could emerge as a byproduct of GCs experiencing the same intrinsic and extrinsic temporal cues as all other cells in the cerebellum. Alternatively, the developmental functions of genes downregulated in GCPs during prenatal development could suggest a reduction in cell fate plasticity and increasing commitment towards the GC fate. Moreover, the timing of GC neurogenesis has been shown to vary across cerebellar lobules (*71*) and to affect the axonal projection patterns of GCs (*72*), but understanding how such differences relate to gene expression and regulation warrants further study.

### Evolutionary dynamics of CREs

We also characterized the evolutionary dynamics of CRE activity across cell types and developmental stages. A limitation of our work is that we can only assess conservation of the regulatory landscape through the conservation of CRE sequences. Comparative studies have shown that regulatory activity, especially for distal elements, typically evolves faster than DNA sequences (*25*, *26*, *28*, *73*). However, here we did not attempt to define regulatory conservation in absolute terms, but only performed comparisons across cell types and developmental stages. To this end, sequence conservation is one of the strongest predictors of regulatory element activity conservation (*25*, *26*, *74*), suggesting that, in the absence of comparative developmental epigenomic atlases at single-cell resolution, differences in sequence constraint can be extended to infer differences in activity conservation.

We detected a common trend across all cell types, with both distal CRE age and sequence conservation decreasing during development. This adds to the accumulating molecular evidence (*21*, *30*, *33*, *75*) in support of von Baer’s observations from the nineteenth century that morphological similarity between developing embryos from different species decreases during development (*76*). This developmental pattern is largely explained by cell type differentiation and maturation, but also by additional temporal differences between cells from matched differentiation states. In both cases, CRE pleiotropy is intertwined with evolutionary constraint (*51*, *69*). Both pleiotropy and regulatory conservation decrease during development and differentiation, in accord with observations in gene expression studies (*30*, *33*).

Besides these shared developmental patterns, we also identified differences in regulatory conservation between cell types. The regulatory landscape of microglia, the immune cells of the brain, shows the fastest evolutionary turnover among all cerebellar cell types. On the other hand, mature astrocytes, including Bergmann glia, have the most conserved distal CREs, not only in the adult cerebellum, but also across a wide selection of cell types in adult mouse organs. Given our observations pertaining to the effect of development and pleiotropy on evolutionary conservation, this difference might be explained by astrocytes maintaining some of the functional properties of neural progenitors, including the ability to reactivate proliferation after injury (*77*), and/or by stronger pleiotropic constraints due to their bridging interactions with multiple cell types of the otherwise disconnected neuronal and vascular networks in the brain (*78*). Finally, among neuron types, throughout development and especially in embryonic stages, GCs are characterized by reduced conservation, which might be associated with the more recent evolutionary emergence of the proliferative external granule cell layer (*6*, *7*) as well as the overall degree of developmental adaptations in GCs across vertebrates (*4*).

Despite these differences between cell types, our data suggest that previous observations regarding declining regulatory constraint during organ development (*21*) are largely explained by changes within cell types during development rather than temporal changes in cellular composition. Given that the cerebellum has been successfully used as a model system to study cell fate specification, neurogenesis and other developmental processes (*4*, *10*, *24*), we expect that many of our observations regarding the developmental and evolutionary dynamics of regulatory elements, and their interplay, are generally applicable to many mammalian organs.

## Acknowledgements

We thank Duncan Odom and Sebastian M. Waszak for critical reading of the manuscript, and Simon Anders, Francesco Lamanna, Florent Murat, Oliver Stegle, Judith Zaugg and members of the Kaessmann lab for discussions. We thank Julia Schmidt and Céline Schneider for technical support.

## Author contributions

H.K., M.C.M., I.S. and M.S. originally designed the study; M.S. collected samples; M.S. and R.F. generated data; I.S. performed all analyses with contributions from M.S. and key input from R.F., K.L. and N.T; E.L., L.M.K., K.O. and P.J. provided important feedback and discussions; H.K. and S.M.P. provided funding; H.K., S.M.P. and M.C.M. supervised the study; I.S., M.S., M.C.M. and H.K. wrote the manuscript with input from all authors.

## Data and code availability

Data generated in this study have been deposited in ArrayExpress with the accession code E-MTAB-9765. All code used to analyze the data is available online at https://github.com/ioansarr/mouse-cerebellum-atac/. Processed data can be interactively explored at https://apps.kaessmannlab.org/mouse_cereb_atac/. Active CREs across cell types and stages (Methods) can be inspected as custom UCSC genome browser tracks at: http://genome.ucsc.edu/s/ioansarr/Mouse%20cerebellum%20snATAC-seq%20cCREs.

## Materials and Methods

### Ethics statement and sample collection

All animal procedures were performed in compliance with international ethical guidelines for the care and use of laboratory animals, and were approved by the local animal welfare authorities (Regierungspräsidium Karlsruhe). RjOrl:SWISS time-mated pregnant mice, litters at postnatal days P0-P14 and adult (P63) mice were purchased from Janvier Labs (France). The adult mice were sacrificed by cervical dislocation, the pups by decapitation. Prenatal samples were collected before noon and are referred to by full completed days post coitum (e.g. E10). Cerebella from animals from both sexes were dissected as whole or in 2 halves (Table S1). E10 and E11 samples were pooled from 2-4 littermates. During dissection the developmental samples were kept in ice-cold PBS and most of the meninges was removed. All samples were snap-frozen in liquid nitrogen.

### Single nucleus ATAC-seq data generation

To isolate nuclei from frozen pre- and postnatal cerebellum samples we made use of a published protocol (*79*), which we optimized with the aim to maximize the recovery of nuclei. The frozen tissue was homogenized on ice in 250 mM sucrose, 25 mM KCl, 5 mM MgCl_2_, 10 mM Tris-HCl (pH 8), 0.1% IGEPAL, 1 μM DTT, 0.4 U/μl Murine RNase Inhibitor (New England BioLabs), 0.2 U/μl SUPERas-In (Ambion), cOmplete Protease Inhibitor Coctail (Roche) by trituration and/or by using a micropestle. After 5 minutes incubation the remaining bits of unlysed tissue were pelleted by centrifugation at 100 g for 1 minute. The cleared homogenate was centrifuged at 400 g for 4 minutes to pellet the nuclei. Nuclei from pre- or postnatal samples were washed in the homogenization buffer once or twice, respectively, collected by centrifugation and resuspended in the Nuclei Buffer (10x Genomics). If needed the nuclei were strained using 40 μm Flowmi strainers (Sigma). For estimation of nuclei concentration, Hoechst DNA dye was added and nuclei were counted on Countess II FL Automated Cell Counter (Thermo Fisher Scientific). 15,000 nuclei per sample were subjected to tagmentation, single cell barcoding and library construction using the Chromium Single Cell ATAC Reagent kits (v1) and Chromium Controller instrument (10x Genomics) according to the manufacturer’s protocols. Libraries were amplified in 10 PCR cycles and quantified on Qubit Fluorometer (Thermo Fisher Scientific). Average fragment size was determined on Fragment Analyzer (Agilent). Libraries were sequenced on Illumina NextSeq 550 (34 cycles for both Read 1 and Read 2, 8 cycles for i7 index and 16 cycles for i5 index) with ~220 million read pairs per library.

### snATAC-seq processing and quality control

Raw sequencing data were demultiplexed and converted to fastq format using cellranger-atac mkfastq (1.1.0) (*36*). The command cellranger-atac count (1.1.0) was used to correct droplet barcodes for sequencing errors, align reads to the mouse genome (mm10), generate position-corrected tabular fragment files and identify PCR duplicates from fragments with identical positions originating from the same droplet barcode. Position-corrected fragments were used as an input to ArchR (0.9.2) for subsequent processing, quality control and analysis (*44*). Using ArchR we tiled the genome in 500 bp bins and counted fragments per barcode and bin. Gene activity scores were inferred by counting all fragments overlapping a gene’s body and applying an exponential decay weighting function to distal regulatory elements. We used a series of established quality control metrics to assess the quality of our libraries. All samples included in our dataset showed the expected periodicity in the frequency of fragment sizes (Fig. S1A) and a minimum 5-fold enrichment of insertions around annotated TSS (Fig. S1B). Following the identification of putative CREs active in cerebellum development (see below) we estimated rank based (Spearman’s) correlations across the accessibility profiles of our samples. For all stages, the highest correlation was observed between biological replicates (Spearman’s rho: 0.94-0.98; Fig. S1E).

### Cell identification and doublet removal

Barcodes corresponding to nuclei of cells were distinguished from empty droplets by requiring at least 5,000 fragments and a minimum TSS enrichment of 3. A total of 117,713 barcodes passed this first filtering step. A common issue of single-cell technologies arises from two or more cells being included in the same droplet and thus sharing the same barcode. Such putative doublets were identified *in silico* using the functions addDoubletScores() from ArchR. This method is based on simulating artificial doublets by adding up the fragment counts of random cell pairs from the same sample. The simulated doublets are co-embedded with the real cells and putative doublets are identified as cells that are consistently found near simulated doublets in the embedding. We used latent semantic indexing (LSI, as described in *“Dimensionality reduction and clustering”*) to embed our cells into 50 dimensions and performed 10 trials, each marking the 10 nearest neighbors of every simulated doublet. Cells from the same sample were then ranked based on their doublet enrichment score, i.e., the number of simulated doublets identified as a nearest neighbor divided by the expected number given a random uniform distribution. We then used the function filterDoublets() to remove the top X% of cells from each sample, where X is proportional to the number of cells per sample, as the fraction of expected doublets depends linearly on the number of loaded cells (*36*). We used a filterRatio of 1, removing 5% from a sample of 5,000 cells (1^st^ pass doublets). 111,201 cells passed this filtering step. However, in downstream stages of the analysis we noticed that cells with high number of fragments and doublet enrichment scores were clustering together. We thus implemented an additional filtering step, removing all cells with more than 45,000 fragments (approximately double the median number of fragments per cell) or a doublet enrichment score greater than 4 (Fig. S1C). This filtering step removed an additional 13,855 cells (2^nd^ pass doublets). We also marked six clusters that were enriched for these putative doublets (Fig. S1D). To assess whether the remaining cells in these clusters also corresponded to doublets or low quality cells, we identified marker genes per cluster using the gene activity matrix as an input to the function getMarkerFeatures() from ArchR. Even after removing the 1^st^ and 2^nd^ pass doublets, four of the six clusters (C5, C18, C21, C55) showed no enrichment for gene activity or a mixture of marker genes for distinct cell types present in the same sample. Thus, we decided to exclude all cells belonging to these four spurious clusters from subsequent analyses. This iterative, stringent filtering procedure resulted in a total of 91,922 high quality cells with a median of 20,558 unique fragments per cell and a median TSS enrichment score of 7.8. Following cell type annotation (see *“Iterative clustering and cell annotation”)* a total of 2,435 cells, some of which might represent remaining doublets, could not be confidently assigned to a cell type and were labeled as “Other/mixture”.

### Dimensionality reduction and clustering

Dimensionality reduction for snATAC-seq data is particularly challenging due to the sparsity of the data. Furthermore, the nearly binary nature of the data does not allow for the identification of highly variable features. To overcome these limitations, we used a recently developed iterative procedure from the ArchR package (0.9.2), based on a widely used method for snATAC-seq data, latent semantic indexing (LSI). LSI scales each feature by the sequencing depth of each cell, as well as with the inverse of its frequency across all cells, thus assigning a higher weight to features with restricted activity across cells (*80*). The transformed features are then used as input for dimensionality reduction based on singular value decomposition (SVD). However, in the absence of additional information about cell types or states, this procedure assigns higher weight to both cell type-restricted CREs, as well as features that show low-level, noisy activity across cells in a cell type-independent manner. The iterative procedure implemented in ArchR overcomes this limitation by performing LSI multiple times in a subset of the dataset. With each LSI round, cells are clustered in the SVD space using Louvain clustering as implemented in Seurat (3.0.1) (*81*), the most variable features between clusters are identified and then used as input for the next LSI round. Thus, with each iteration, ArchR converges towards a set of features that is most informative in distinguishing cell types and states in the dataset.

For our analyses, we used five iterations, each time sampling 20,000 cells (10,000 for subclustering analyses, see *“Iterative clustering and cell type annotation”*) with ten random starts and gradually increasing the clustering resolution (0.1, 0.2, 0.4, 0.8). We used 500 bp-wide genome tiles as input to avoid biases against rare cell types and states, excluded mitochondrial and sex chromosomes and binarized the count matrix. Features were transformed with TF-log(IDF) and the 100,000 most variable features (50,000 for subclustering analyses) were reduced to 100 dimensions with SVD (50 for subclustering analyses). Each component was weighted by the percentage of variance it explained. Components showing an unusually high correlation with sequencing depth (Pearson’s *r* > 0.75) were excluded from downstream analyses. These SVD components were used as input for Louvain clustering and projection to a UMAP embedding (Fig. 1B-C and S1F).

### Iterative clustering and cell type annotation

The multidimensional nature of our dataset poises challenges to cell type annotation as similarity between cells might be driven by factors other than cell type identity, such as developmental stage, or differentiation and maturation state. To overcome this challenge, we implemented an iterative clustering approach to identify high quality cells and assign them to cerebellar cell types and states. First we applied Louvain clustering to the full dataset, identifying 47 distinct clusters (resolution 1.5; Fig. S1F). We used gene activity as a proxy for gene expression to assign clusters to putative cell types and states (Fig. 1E, S2A) based on previously described marker genes (*2*, *13*), as well as manual investigation of *in situ* hybridization data from the Allen Developing Mouse Brain Atlas resource (*58*). We observed that many of our clusters contained additional substructure, which we were unable to recover when considering the entire dataset, even when increasing the clustering resolution. Thus, we decided to subcluster cells from clusters where additional structure was apparent (Fig. S1F-H). We grouped these cells into three major groups (astroglia, early-born neurons, and GABAergic neuroblasts from E13) and repeated the LSI dimensionality reduction and clustering (38, 37 and 21 subclusters obtained with a resolution of 2.5, 2.5 and 2.0 respectively; Fig. S1I-H). We followed the same approach, relying on gene activity of marker genes to identify cell types and their subtypes. Cluster 3 from the astroglia group was further split into three subclusters. We then assigned each cell to the most specific cell type and state label available from the full dataset or subclustering analysis. Following inspection of individual developmental stages (Fig. S2B), we identified a total of 2,435 cells (E11: C13, E12: C14-C15, P14: C12) that could not be confidently assigned to a specific cell type and that likely include remaining doublets or unresolved clusters. These were labeled as “Other/mixture” and excluded from further analyses. Taken together, we were able to provide broad cell type and subtype/cell state labels to 97% of the cells in the dataset.

### Integration with scRNA-seq data

We used single-cell RNA-seq data from a previous study profiling mouse hindbrain and cerebellum development (*12*) to validate our cell type annotation and assess the value of gene activity as a proxy for gene expression. To improve the computational efficiency and accuracy of our approach we performed the integration in a stage-wise manner, matching developmental stages between studies as closely as possible. We reprocessed the RNA-seq data per stage using a widely-used pipeline as implemented in the package Seurat (3.0.1) (*81*). Briefly, we used high quality cells, as previously filtered by the authors, applied SCTransform() to scale the data and identify highly variable genes, and reduced the dimensions of the data to 50 principal components.

We also reprocessed our snATAC-seq data in a stage-wise manner using ArchR as described above (see *“Dimensionality reduction and clustering”*; Fig. S2B) and with the following modifications: 50,000 variable features, 50 SVD components, clustering resolution of 1.0. When analyzing the data in a stage-wise manner, we observed mild batch effects in the P14 and P63 datasets. Given the very high correlation between the biological replicates (Spearman’s rho: 0.98 for both stages), we reasoned that the separation between the batches was more obvious due to the low cellular complexity of these samples, which are both dominated by GCs. We corrected these mild batch effects using Harmony (1.0) in the LSI space (*82*). Although batch effects were only visible in P14 and P63, we applied the Harmony correction to all stages for consistency in our analysis. We then created a custom Seurat object containing the gene activity scores, metadata and embeddings of each snATAC-seq sample.

The integration between matching scRNA-seq and snATAC-seq datasets was performed using canonical correlation analysis (CCA) as implemented in Seurat (*81*). RNA UMI counts and ATAC-seq derived gene activity scores were scaled down to 10,000 counts per cell and the highly variable genes from the RNA-seq sample (reference) for which gene activity estimates were available in the ATAC-seq sample (query) were used as features to identify transfer anchors in the CCA space. We then transferred cell type labels and RNA-seq UMI counts to the snATAC-seq Seurat object using the 10 (for cell type labels) and 15 (for RNA imputation) nearest neighbors in the LSI space to weigh the predictions. We estimated the Jaccard similarity index between our cell type annotations and those transferred from the RNA-seq data and visualized the results in a heatmap (only showing sets that had a similarity index of at least 0.15 with another group).

Overall, we observed high concordance between the two annotations (Fig. S2C). However, there were three notable discrepancies. First, GABAergic DNs in our dataset best matched a scRNA-seq cluster annotated as excitatory (glutamatergic) nuclei (DN) neurons. GABAergic and glutamatergic DNs share several marker genes and are both abundant in early cerebellum development (Fig. 1 and S2A). Given that GABAergic DNs were not identified as a distinct cell type in the scRNA-seq data, we hypothesized that the scRNA-seq cluster could contain a mixture of both glutamatergic and GABAergic DN neurons. Indeed, we observed additional substructure in this scRNA-seq cluster with the majority of the cells expressing *Gad2* and *Sox14* (Fig. S2F), which are markers for GABAergic DNs (*83*). Second, our differentiating GCs during prenatal development were labeled as unipolar brush cells (UBCs) after the integration with the scRNA-seq data. As before, we discovered additional substructure within this cluster (Fig. S2E), with only a fraction being positive for the UBC marker *Lmx1a* (*84*), and a sizeable group of cells expressing the GC-specific marker *Kcnd2*. Third, differentiating PCs in our dataset matched with brainstem progenitors in the scRNA-seq atlas. However, given that we dissected cerebella, we expect only small contaminating contributions from the brainstem. Indeed, we identified a small population of brainstem progenitors in our data (222 cells, 80% of which from E10-E12), characterized by high gene activity for several Hox genes (*Hoxc13, Hoxc6, Hoxc4, Hoxa3*). These are expressed in the lower brainstem (pons and medulla) but not in the cerebellum (*4*). Our PC differentiating population showed no activity for these Hox genes and was instead enriched for PC markers, such as *Skor2* and *Lhx5* (Fig. S2A).

Compared with our study, the scRNA-seq dataset includes entire hindbrain samples for E10-E12, cerebellum samples from shifted stages for later prenatal development and no P63 samples. Despite these differences, we observed high concordance in the cell type annotations (Fig. S2C) and cell type prediction scores (median prediction score: 0.81). Similarly the imputed RNA-seq data of the genes used for the integration were robustly correlated with the gene activity scores obtained from the snATAC-seq data (median Pearson’s *r*: 0.5; Fig. S2D). This high correlation motivated us to use gene activity scores as a proxy for gene expression for downstream analyses. We preferred gene activity to imputed RNA expression because it was derived directly from our data, thus overcoming issues with the compatibility of the dissections and developmental stages between the two studies.

### Identification and annotation of putative CREs

#### Cluster and sample-based pseudobulk generation and peak calling

We used ArchR’s function addReproduciblePeakSet() to identify regions of accessible chromatin, as a proxy for putative CREs (Fig. S3A). We opted for this framework because it allowed us to account for differences in the abundance and heterogeneity of cell types in our dataset and to utilize our biological replicates to assess peak reproducibility. Briefly, we used the 47 clusters from the full dataset to generate pseudobulks comprised of cells from the same cluster and sample (biological replicate). This allowed us to identify cell type/state-specific peaks while retaining the information of the sample the cells came from (and thus filtering out irreproducible peaks). Depending on the number of cells contained in each sample, we constructed 2-10 pseudobulk groups, each containing a minimum of 100 and a maximum of 1,000 cells, while limiting the maximum number of fragments per sample to 50 million. The minimum threshold for the number of groups and cells (2 x 100 = 200) was estimated by subsampling analysis as the minimum number of sampled cells required to obtain a Pearson’s correlation of at least *r*=0.90 with the entire pseudobulk of a given cluster. The rationale behind setting a maximum threshold was to limit the contribution of very abundant cell types and states (e.g., mature GCs) to the peak set. We chose to restrict this parameter primarily through the number of cells in a group (1,000) rather than the number of samples, to facilitate the identification of CREs from cell types that are consistently rare in our data but present in multiple samples (e.g., microglia or postnatal PCs). If a cluster did not have 100 cells from at least two samples, we allowed up to 80% of the cells from a group to be resampled with replacement to reach the required number. In practice, this only affected two clusters (C1: erythroid and C15: mature UBCs). We then called peaks within each group (cluster + sample) using MACS2 (2.1.2) (*85*) through ArchR with the following parameters “--shift −75 --extsize 150 --nomodel --call-summits --nolambda --keep-dup all -q 0.01”. We allowed up to 200,000 peaks to be identified from a single group, or up to 1,000 peaks per cell (thus for a group of 50 cells that would be 50,000 peaks).

#### Union peak set generation

Calling peaks in a cluster- and sample-aware manner resulted in multiple peak sets that had to be collapsed into a single peak annotation. Given that many of our subsequent analyses depended on the overlap with various genomic elements (see *Peak annotation*), which can be biased by the width of each peak, we followed ArchR’s iterative overlap merging procedure to obtain the maximum number of 500 bp-wide non-overlapping peaks (Fig. S3A). Briefly, ArchR’s approach is based on extending peak summits from each group by 250 bp in each flank, then identifying overlapping windows and ranking them based on peak’s significance, normalized by the sequencing depth of each group. The most significant peak is retained and all other overlapping peaks are removed. This approach avoids chain-merging of adjacent peaks as well as the removal of peaks that are proximal but non-overlapping to the most significant peak (*44*). For some analyses we still allowed for the presence of broader peaks by identifying streaks of adjacent peaks with very similar activity patterns throughout the dataset and grouping them into a shared functional unit (see “*Peak co-accessibility and broad CRE identification”*). We required each peak to be supported by at least two biological samples (i.e., replicates), which resulted to 499,146 non-redundant peaks (permissive peak set). To avoid the inclusion of noisy peaks from clusters with very large numbers of cells or replicates, we further filtered for peaks that were active in at least 5% of at least one cluster (robust peak set; Fig. S3A-B). These robust peaks were associated with higher summit scores and reproducibility and were enriched for overlap with H3K27ac histone marks during hindbrain development and were more likely to be assigned to putative target genes (see *Assignment to putative target genes*). Thus, reasoning that these robust peaks were more likely to correspond to putative CREs, we limited all our downstream analyses to this set. However, we also provide the permissive set as a resource as it could contain CREs with more restrained activity to specific cell types/states (https://apps.kaessmannlab.org/mouse_cereb_atac/).

#### Peak annotation

ArchR automatically annotated peaks based on their genomic context as promoters (−2,000/+100 bp from a TSS), exonic, intronic or intergenic based on UCSC annotations for the mm10 genome (*86*). We integrated this annotation with the biotypes of the associated gene, acquired from Ensembl Biomart (v94) (*87*), and collapsed to four main biotypes: protein-coding, lncRNA (including antisense, bidirectional and intergenic lncRNAs), small RNA (miRNA, snRNA and snoRNA) and other (mostly pseudogenes). We further supplemented the UCSC annotation with a recently described set of lncRNAs expressed during mouse organ development (*33*). We used bedtools intersect (2.29) (*88*) to identify overlaps between peaks and lncRNA promoters, exons and gene models, updating the annotations of previously intergenic and intronic CREs based on their overlap with lncRNA features.

To determine peak overlap with TF motifs and to assess TF activity based on the aggregate accessibility of these motifs, we used the ArchR implementation of the package chromVAR (*66*). We scanned our peak set for instances of the motifs present in the cisBP collection (*89*) using the default cutoff of *P* < 5e-05. We then computed per-cell deviations for all motifs using a GC-content matched background peak set.

To assess sequence constraint in putative CREs we used estimates of evolutionary conservation (phastCons scores) based on multiple alignments (*60*). We downloaded phastCons scores for vertebrates and eutherian mammals (commonly referred to as placental mammals) for mm10 as bigWig files from the UCSC table browser. As selective constraint is often concentrated to specific regions of a CRE (e.g., TF binding sites), we employed a sliding window approach to identify the most conserved 100 bp region of each CRE using the UCSC utility bigWigAverageOverBed (*90*). The mean phastCons score within the most conserved 100 bp region was considered as a metric of constraint for each CRE.

To assign a minimum evolutionary age for each putative CRE we used syntenic alignments between mouse and 16 high quality vertebrate genomes of various phylogenetic distances (Fig. S6A, Table S5) to assess the presence or absence of our putative CREs in the genomes of other species. We downloaded chain alignments from UCSC and used liftover (-minMatch=0.1 -multiple -minSizeQ=50 -minSizeT=50) to identify syntenic regions in each species. We then assigned a putative age to each CRE based on the estimated age of divergence between mouse and the most distant species in which a syntenic region could be detected. We observed that older regions were also found in more species, suggesting that the number of false positive alignments was overall low (Fig. S6B).

To determine the contribution of transposable elements to the identified putative CREs, we used bedtools (2.29) (*88*) to intersect our CREs with the RepeatMasker track for the mm10 genome, downloaded from UCSC Table Browser (*91*). Per CRE, we estimated the fraction covered by repeats and retained the name and class of the overlapping repeats.

#### Peak co-accessibility and broad CRE identification

Imposing a fixed width to our peaks was favorable for many downstream analyses but precluded us from identifying broad regions of accessible chromatin, such as super-enhancers, which have been reported as important regulators of cell identity (*92*, *93*). To facilitate the identification and characterization of such elements during mouse cerebellum development, we grouped adjacent peaks with correlated accessibility profiles into broad CRE units (Fig. S3A). First, we computed correlations in accessibility profiles between peaks located up to 100 kb from each other using ArchR’s addCoAccessibility() function. Since the sparsity of single-cell data can preclude the identification of correlations, we created 5,000 pseudocells (groups of 50 nearest neighbors based on their distance in the LSI space) and aggregated their accessibility profiles. Pseudocells showing more than 50% overlap in their members with another aggregate were excluded resulting to a total of 4,083 groupings. The aggregated accessibility profiles were log normalized and pair-wise Pearson’s correlations were computed between neighboring peaks. Focusing on directly adjacent peaks, we observed a strong negative correlation between peak distance and co-accessibility with a clear elbow point at 1,000 bp and a correlation coefficient of 0.4 (data not shown). Thus, we grouped together peaks with a Pearson’s *r* of at least 0.4 and a maximum distance of 1,000 bp into a common regulatory unit. We allowed consecutive chains of adjacent and co-accessible peaks to be grouped together into a single regulatory unit. The maximum number of peaks in a chain was 35 but 75% of the chains contained up to 3 peaks.

#### Assignment to putative target genes

As CREs can act over larger distances in the linear genome, the identification of their target genes is particularly challenging. Whereas *bona fide* regulatory interactions can only be established by perturbation or genetic studies, CREs and their target genes are expected to be active in the same cell types/states. We thus used the correlation between peak accessibility and gene expression (approximated by gene activity scores) across cell types and states in our dataset to assign CREs to their putative target genes, akin to previous studies (*22*, *39*). We opted for using gene activity over imputed gene expression from the integration with the scRNA-seq data (see *“Integration with scRNA-seq data”*), because matched transcriptomics data were not available for some stages due to differences in dissections and sampling, whereas the two metrics were overall highly correlated in properly matched stages (Fig. S2D). We used the same cell groupings with the peak co-accessibility analysis (see *Peak co-accessibility and broad CRE identification*) and ArchR’s function addPeak2GeneLinks() to compute Pearson’s correlations between gene activity and the accessibility of all peaks within 250 bp upstream or downstream of the gene’s TSS. Although ArchR provides parametric estimates of significance for these correlations, we observed that the obtained *P*-values were very small even for low correlation coefficients, likely due to the large size of our dataset. We thus adapted a previously described method based on computing interchromosomal correlations to obtain an empirical null distribution and identify a biologically meaningful correlation cutoff (*94*). We randomly selected 1,000 peaks from chromosome 1 and correlated their expression with 1,000 random genes from chromosomes 2, 3 and 4. The 99^th^ quantile of these interchromosomal correlations was *r*=0.425. Thus, we identified peak-to-gene links based on a cutoff of 0.45 (FDR < 1%; Fig. 2B). To estimate the number of distal CREs associated with each gene (Fig. 2C), we collapsed CREs to their broad regulatory units *(see “Peak co-accessibility and broad CRE identification”)* and counted the number of non-redundant (i.e., unique or belonging to different broad regulatory units) distal CREs per gene.

#### Benchmarking of putative cerebellum CREs with external datasets

We compared the identified putative CREs with relevant bulk and single-cell datasets from previous studies. We downloaded Encode chromHMM annotations (*50*) for mouse candidate CREs for a variety of different tissues and developmental stages (Table S5) and intersected them with our peaks using bedtools (2.29) (*88*). For each sample (tissue and developmental stage) we estimated the fraction of Encode-annotated enhancers (“Enh”) and heterochromatin (“Het”) elements that overlapped our peaks. Predicted enhances active in the nervous tissues and in particular hindbrain showed very high overlap with our peaks, whereas the percentage of recovered enhancers for tissues of mesoderm and endoderm origin was much lower (Fig. S3C). The same was true for experimentally validated enhancers (Fig. S3E) (*49*). By contrast, our peaks were depleted of putative heterochromatin regions for the nervous tissues, whereas the overlap was higher, although overall below 20%, for the other tissues (Fig. S3D). We also benchmarked our peak set against a recently published single-cell atlas of chromatin accessibility in adult mouse organs (*38*). After identifying overlapping peaks using bedtools intersect, we estimated the fraction of reads per cell in the mouse atlas within regions overlapping our peak set. Cells of cerebellar origin showed the highest fraction of reads in our peaks (median 80%; Fig. S3F).

#### Characterization of enriched CRE sets

Throughout this study we aimed to characterize groups of CREs with similar activity patterns (e.g., belonging to the same cluster, showing differential accessibility across developmental stages or during differentiation). To associate CREs with the biological processes (BP) of their putative target genes we used the R implementation of GREAT (rGREAT, 1.18.0) (*95*) to identify enriched gene ontologies. Similarly, we used Homer (4.11.1) (*96*) to identify enriched TF motifs in each cluster of CREs. We performed motif search using the command findMotifsGenome.pl with the options –gc to account for GC content biases and –size given to scan the entire peak regions. For both BP and motif enrichment all CREs tested in the analysis were combined and used as a background peak set. The *P*-values of the enrichment of each term for each CRE set were capped for visualization purposes (typically to 10^−100^ for motifs and 10^−30^ for BP enrichment) and plotted in a heat map. For BP enrichment, when the number of CRE sets was large and/or processes were too similar, we collapsed BP terms to a non-redundant set using Revigo (*97*) with the following parameters: allowed similarity: 0.5, semantic similarity measure: SimRel.

### CRE activity across cell types and developmental stages

To profile the activity of CREs during the development of different cerebellar cell types, we aggregated the activity of all cells into pseudobulks for a given cell type and developmental time point. We excluded samples from cell type mixtures (e.g., NTZ, Parab. and isth. N., Other/mixture) and non-neural cell types (e.g., erythroid, vascular), as well as pseudobulks comprised of fewer than 50 cells. We only considered robust CREs for this analysis. We estimated CPM values to account for differences in cell numbers and sequencing depth and standardized our data by scaling by the maximum CPM value of each CRE.

We used a two-stage clustering approach inspired by a recent study (*98*). First we performed K-means clustering with a high number of primary clusters (n=50). Then, for each primary cluster we estimated the mean activity pattern and used it as input for hierarchical clustering (based on the correlation distance matrix between cluster centers). We then used the clustering dendrogram to iteratively merge the two most similar branches and compute the silhouette score for each new set of clusters. We determined the optimal number of final clusters to be 26, based on the distances between the hierarchical clustering branches, the silhouette score distribution of each iterative merge, and the original number of optimal clusters suggested by the silhouettes of different K-means runs.

We further assessed the clustering quality for each CRE by estimating the correlation between its activity and its cluster’s mean. The median Pearson’s *r* estimate for a CRE with its own cluster was 0.63. CREs with a correlation estimate of at least 0.5 with their cluster center were considered to be “confident cluster members”. These “confident cluster members” were associated with gene ontology terms based on their nearby genes using GREAT and enriched TF motifs using Homer (see *Characterization of enriched peak sets*). To visualize the clustering results (Fig. 2D), we randomly selected 50,000 CREs (with *r* > 0.5 to their cluster mean or belonging to the ubiquitously active cluster) and plotted their standardized activity across cell types and developmental stages in a heat map. We chose random selection over prioritizing highly variable features to better represent the true proportions of CREs in each cluster.

To generate custom tracks of cell type- and time-specific CRE activity for the UCSC genome browser (http://genome.ucsc.edu/s/ioansarr/Mouse%20cerebellum%20snATAC-seq%20cCREs), we used the same pseudobulks generated for the clustering analysis. We selected active CREs per cell type and developmental stage as those with CPM ≥ 5 and exported them in bed format. Bed files per cell type and stage were subsequently uploaded to the UCSC browser.

### Periods of greater change in chromatin accessibility

To identify periods of greater regulatory change we aggregated the accessibility profiles of all cells per cell type, developmental stage and individual (biological replicate). We only consider the most abundant cell types in our dataset (Astroglia, GABA DN, Glut. DN, PC, INs, GC) and samples with at least 1 million fragments per replicate and cell type, which we determined as the minimum number of fragments to detect any differentially accessible regions (Fig. S4B). Only autosomal CREs were included for this analysis as our biological replicates are from different sexes. For each cell type, we used DESeq2 (1.26) (*99*) to identify differentially accessible CREs across adjacent developmental stages requiring a Benjamini-Hochberg adjusted *P*-value < 0.05 and an absolute log_2_ fold change >= 0.5. The overlap between differentially accessible CREs over multiple stages and cell types was visualized using the R packages VennDiagramm (*100*) and UpSetR (*101*).

To assess the effect of library size on our ability to identify differentially accessible regions, we performed a downsampling analysis in the comparison of GCs between P14 and P63. Although 4 million fragments were sufficient to achieve high correlation (Spearman’s rho > 0.95) with the ground truth (full pseudobulk), the number of identified differentially accessible CREs increased with larger library sizes (Fig. S4B). To control for differences in library size between cell types and developmental stages, we performed the same analysis described above (Fig. 3B), downsampling all pseudobulk libraries to 4 million fragments, excluding samples that had fewer fragments. Although the number of detected differentially accessible CREs decreased, greater regulatory change was detected in the same developmental windows (Fig. S4C).

Low cell numbers decrease the power to detect differentially accessible regions (as a metric of dissimilarity), but increase dissimilarity estimated by correlations due to the introduction of stochastic noise (Fig. S4B). We thus decided to use correlations across adjacent stages as an orthogonal approach to validate the periods of greater regulatory change. We aggregated CRE accessibility profiles across cells from the same cell type and developmental stage and estimated log_10_-transformed CPM values. For each cell type we calculated Pearson’s correlations across adjacent stages for the developmental windows considered in Fig. 3A. We chose Pearson’s over ranked-based correlations to allow for a larger contribution of radical changes in CRE accessibility towards the dissimilarity estimate.

For the quantification of cellular composition changes, we estimated the fraction of cells belonging to each cell type per sample (developmental stage and biological replicate). To compare the cellular composition across samples, we calculated absolute changes in the fraction of each cell type and then computed their sum. Thus for each pairwise comparison we estimated a score ranging from 0 (identical composition) to 2 (completely different cell types in the two samples). To assess the significance of temporal differences while accounting for biological and technical confounders, such as differences across individuals, tissue dissection, nuclei preparation and *in silico* filtering, we compared differences in cell type proportions between samples from adjacent developmental stages and biological replicates from the same developmental stage using a t-test (Fig. 3E).

### Cell type differentiation

For each of the three major cerebellar neuron types (GCs, PCs and INs), we identified cells assigned to that cell type and reanalyzed them separately. We only considered developmental stages with at least 100 cells assigned to the specified cell type. We applied the IDF transformation to robust CREs accessible in at least 1% of the cells and embedded the data into 20 components using SVD. In agreement with previous studies (*38*), the first SVD component showed very high correlation with the sequencing depth of each cell and was therefore excluded from downstream analyses. We chose a single-step IDF-SVD analysis over an iterative LSI (see *“Dimensionality reduction and clustering”*), because the latter, although useful for the identification of distinct cell types in the full dataset, exaggerates differences between cells by prioritizing CREs with variable accessibility across clusters (the borders of which are arbitrary in developmental or spatial gradients). We also observed that performing the IDF transformation without scaling by each cell’s sequencing depth (TF) and omitting the 1^st^ SVD outperformed the standard TF-IDF in capturing continuous differentiation signals (inferred based on the activity of known marker genes).

As we observed developmental signal that was orthogonal to differentiation (e.g., pre- and postnatal GCPs and GC neuroblasts), these components were further corrected using Harmony (1.0), a single-cell integration algorithm that relies on soft-clustering, thus allowing for smooth transitions between cell states, such as those observed during cell differentiation (*82*) (Fig. S7A-C). To allow for differentiation states unique to individual developmental stages (e.g., PC neuroblasts in E12) we set the ridge regression penalty parameter to a relatively large value (lambda=5). The obtained Harmony-corrected dimensions were then used as input for clustering and UMAP projection.

We used diffusion-based pseudotime (*64*), as implemented in scanpy (1.4.5) (*102*) to approximate differentiation and maturation processes. After transferring the data to a scanpy object, we used the Harmony-corrected components to construct a graph based on the 20 nearest neighbors of each cell. A diffusion map was computed based on the neighborhood graph and was used as input for pseudotime estimated with zero branchings. The root of the pseudotime was specified as a random cell belonging to the earliest timepoint and from a cluster characterized by the expression of precursor marker genes for the respective cell type (e.g., *Atoh1* in GCs, *Ptf1a* in INs). Specifically for the INs, where multiple subtypes are specified at different developmental stages (Fig. S7D), we observed that pseudotime values above 0.6 captured subtype rather than differentiation signal. Since the aim of this analysis was to describe differentiation paths common to all IN subtypes, we capped pseudotime values to 0.6, and then rescaled them from 0 to 1 (Fig. S7D).

To identify dynamic features, pseudotime values were divided into 50 bins based on their ranks and cells belonging to the same bin were grouped together. The CRE accessibility, gene activity and chromVar-derived motif deviations profiles were aggregated across cells within each pseudotime bin. Dynamic features across pseudotime were identified based on mutual information between the feature (scaled CRE accessibility, gene activity, chromVar deviation score) and the mean pseudotime of each bin using the function cminjk.pw() from the R package mpmi (*103*). Cutoffs for mututal information index (MMI) were determined based on empirical null distributions obtained by shuffling the pseudotime values across bins (controlling for an FDR < 1%). Furthermore, as the MMI distribution was often bimodal, we used a Gaussian mixture model, as implemented in the R package mclust (5.4.5) (*104*) to split the distribution into two groups. Features belonging to the group with the highest MMI and passing the 1% FDR cutoff were identified as significantly dynamic. Significantly dynamic CREs were Z-score standardized and clustered based on their pseudotemporal profiles using a fuzzy c-means clustering algorithm for time series from the R package Mfuzz (2.46.0) (*105*). The optimal number of clusters was determined by generating an elbow plot for the minimum distance between cluster centroids, as well as by monitoring the trajectory profile and gene ontology enrichment of each cluster (reducing the number of clusters when multiple clusters shared similar trajectories and gene ontology enrichment).

For the integrative analysis of neuronal differentiation we determined overlaps between dynamic CREs across neuron types and visualized them with a chord diagram from the R package circlize (*106*). The significance of the overlap between clusters from different cell types was assessed with hypergeometric tests. For the PCA analysis, we generated pseudobulks per cell type and pseudotime bin, applied a variance stabilizing transformation as implemented in the package DESeq2 (1.26) (*99*) and performed PCA as implemented in the package FactoMineR (*107*).

### Transcriptional regulators of cell type differentiation

The aggregated accessibility of CREs containing a given TF motif can be used to infer the activity of the TF in a given cell type and state. Compared with assessing the activity of the TF using gene expression data (RNA-seq), this metric allows the detection of changes in activity driven by posttranslational modifications, such as phosphorylation or interactions with co-factors. However, TFs that belong to the same family often recognize very similar motifs, hindering the precise identification of the specific TF that is driving the observed change in chromatin accessibility. To overcome this limitation we used an integrative approach to identify TFs with dynamic motif accessibility and gene activity (as a proxy for gene expression) across cell type differentiation. We identified significantly dynamic motifs and gene activity scores based on the MMI estimated from the 50 pseudotime bins (see “*Cell type differentiation”*). We further estimated Pearson’s correlations between motif activity and gene scores across pseudotime (using the average values in the 50 bins). We then identified candidates for the regulation of cell type differentiation as TFs that 1) reached a minimum gene activity score of 1 in at least one pseudotime bin, 2) showed dynamic chromVar deviation score and 3) gene score, and 4) showed an absolute correlation coefficient of *r* > 0.6 between the motif accessibility and gene activity. TFs were then classified as putative activators or repressors depending on the sign of the correlation coefficient between gene activity and motif accessibility akin to similar analyses by others (*67*). Information on TF families was obtained from cisBP (Build 2.0) (*89*).

### Analysis of cerebellar progenitors

The embedding presented in Fig. 4 was generated as described in the section *“Cell type differentiation”*. The identification of progenitor types was performed using the iterative clustering procedure in the entire dataset as discussed in *“Iterative clustering and cell type annotation”*. Marker genes per progenitor type and developmental stage were identified using the function getMarkerFeatures() from ArchR, filtering for FDR < 0.01 and log_2_ fold-change > 1. For plotting, marker genes were prioritized based on the product of their log_2_ fold-change estimate and the –log_10_ *P*-value of the test.

We aggregated chromatin accessibility across cells from the same progenitor type and developmental stage and scaled to a total of 10^6^ fragments per group. Only pseudobulks with at least 100 cells and robust CREs that reached at least 5 CPM in at least one pseudobulk (122,572 putative CREs) were considered for subsequent analyses. For the clustering across progenitor types and stages, we calculated Spearman’s correlation coefficients across samples and performed hierarchical clustering using ward.D2 and pairwise complete observations. To assess the robustness of the clustering we used bootstrapping as implemented in the package pvclust (*108*) with 1,000 repetitions. To identify major patterns of CRE activity in progenitor cells we scaled the 122,572 putative CREs by their maximum activity across the progenitor samples and then used the clustering approach described in *“CRE activity across cell types and developmental stages”*. For this analysis we generated 30 clusters with k-means clustering, which we then collapsed to a final number of 12 clusters based on hierarchical clustering of their cluster means.

For the analysis focusing on the relationship between anterior VZ and bipotent progenitors, we first extracted progenitors from the developmental stages E13-E15 and used ArchR’s getMarkerFeatures() to identify CREs that are specific to each germinal zone, filtering for FDR < 0.01 and log_2_ fold-change > 1. These led to the identification of 1,753 CREs, 238 of which were specific to the bipotent progenitors. We then estimated the fraction of fragments per cell that overlap those CREs and compared early (E10-E12) progenitor populations of different germinal zones. As a complementary analysis, we computed Spearman’s correlations between bipotent progenitors from E13 and E15, and early (E10-E12) progenitor populations based on their entire chromatin accessibility profiles (in the 122,572 putative CREs considered for this analysis).

### CRE evolutionary dynamics

To compare CRE evolutionary dynamics across cell types and developmental stages, we calculated summary statistics per cell based on all active CREs. As described above (see *“Peak annotation”*), for each CRE we inferred its minimum age, estimated the average sequence constraint in its most conserved 100 bp region, and calculated the fraction of the CRE covered by genomic repeats. For each cell, we calculated the mean values for these estimates across all distal (intergenic and intronic) CREs that were accessible in that cell. We focused these analyses on distal elements, to avoid biases from overlaps with protein-coding sequences, which overall show very high sequence conservation, often independently of regulatory constraint. Due to the sparsity of snATAC-seq data, these estimates vary across single cells, even within the same cell type and state. To estimate summary statistics for groups of cells (e.g., as shown in Fig. 5), we estimated 95% confidence intervals using the function CI() from the R package Rmisc (*109*) and only considered estimates based on at least 50 cells.

For the contributions of different age groups and of TE classes to the regulatory landscape of different cell types we estimated the fraction of fragments in CREs in each class (inferred age or overlapping TE class) over the total number of fragments in CREs per cell. The absolute value of this estimate depends on the overall abundance of such elements in the mouse genome (e.g., fraction of fragments in B1 elements is much higher than ERVs), but we observed additional variation between cell types and stages (Fig. 5F and S6H). For the comparison between CREs associated with TFs versus other genes, we considered protein-coding genes with dynamic expression in the developing cerebellum (*30*). Mouse TFs were obtained from AnimalTFDB v3 (*110*). The significance of these comparisons between groups was assessed with two-sided Mann-Whitney *U* tests.

### Temporal differences in GC differentiation

The additive contribution of differentiation (inferred by pseudotime) and developmental stage towards the regulatory landscape of GCs was assessed using a linear model with both pseudotime and developmental stage as predictors of average CRE constraint per cell (as described in *“CRE evolutionary dynamics”*). The significance of each predictor term was estimated using ANOVA tests between the full model and an alternative model that only included the other term.

To detect CREs with radical temporal differences across matched stages of GC differentiation, we first divided cells into five major bins based on their pseudotime values. Biologically meaningful pseudotime cutoffs were identified on the basis of the distribution of pseudotime values (Fig. 7B) and based on the inflexion points of the clusters of CREs that are dynamic during GC differentiation (Fig. 6F). We approximately matched pseudotime bins into differentiation states by identifying marker genes (FDR < 0.01 and log_2_ fold-change > 1) and exploring their expression in the Allen Developing Mouse Brain Atlas resource (*58*). We then examined the distribution of pseudotime values for each developmental stage and bin. We only identified differentially accessible CREs across adjacent developmental stages in the same pseudotime bin when there was no significant difference in the pseudotime value distributions (Fig. S6K) and when all samples (pseudotime bin, developmental stage and biological replicate) had at least 50 cells. For example, we performed no comparisons in pseudotime bin 4, as this stage was highly dynamic and we observed significant differences in the pseudotime values across developmental stages, which confounded temporal and differentiation signals. For comparisons that met these criteria, we used DESeq2 (1.26) (*99*) to identify differentially accessible CREs across adjacent developmental stages requiring a Benjamini-Hochberg adjusted *P*-value < 0.05 and an absolute log_2_ fold change >= 0.5.

For temporal differences in the prenatal GCPs we extracted cells from stages E13-P0 that belonged to the same cluster (cluster 4), prior to any Harmony-correction across developmental stages or pseudotime inference (Fig. 7C and S7A left). Thus, in this analysis we considered cells that overall look the most similar to each other based on their raw chromatin accessibility profiles. We generated pseudobulks per sample (developmental stage and biological replicate), considered CREs with at least 10 counts in at least two samples and applied the variance stabilizing transformation as implemented in DESeq2 (1.26) (*99*). We then detected temporally dynamic CREs using masigPro (1.58.0) with a second degree polynomial and requiring a Benjamini-Hochberg adjusted *P*-value < 0.05 and a goodness of fit value R^2^> 0.7 (*111*). For temporally dynamic CREs, we estimated the mean accessibility across replicates, performed a Z-score standardization across developmental stages and applied soft clustering with k=2 using Mfuzz (2.46.0) (*105*). We examined the activity of each cluster in the rest of the dataset by estimating the fraction of CPM in these CREs per cell. To assess the impact of these temporal differences on gene expression, we considered the genes that were closest to these CREs, excluding genes that were associated with CREs from both clusters. We then examined the expression of these genes in a scRNA-seq atlas of mouse cerebellum development (*12*) focusing on the cells annotated as “Embyronic and postnatal GCPs-1”. We used proximity instead of our correlation-based gene assignment approach to avoid the circularity of examining the concordance between gene expression and chromatin accessibility after requiring them to be correlated (indeed they both show the same temporal pattern in agreement with our proximity analysis; data not shown).

### General statistics and visualization

All statistical analyses were performed in R (3.6.3) (*112*) using the packages tidyverse (1.3.0) (*113*), data.table (1.12.8) (*114*), Matrix (1.2.18) (*115*), SummarizedExperiment (1.16.1) (*116*), irlba (2.3.3) (*117*) and cluster (2.1.0) (*118*). Heatmaps were drawn using the packages ComplexHeatmap (2.2.0) (*119*) and pheatmap (1.0.12) (*120*). Additionally, we used the R packages shiny (1.4.0.2) (*121*), shinyjs (1.1) (*122*), Gviz (1.30.3) (*123*) and GenomicInteractions (1.20.3) (*124*) to build the app that facilitates the interactive exploration of our data.

## Supplementary figures (S1-S7)

**Fig. S1.**
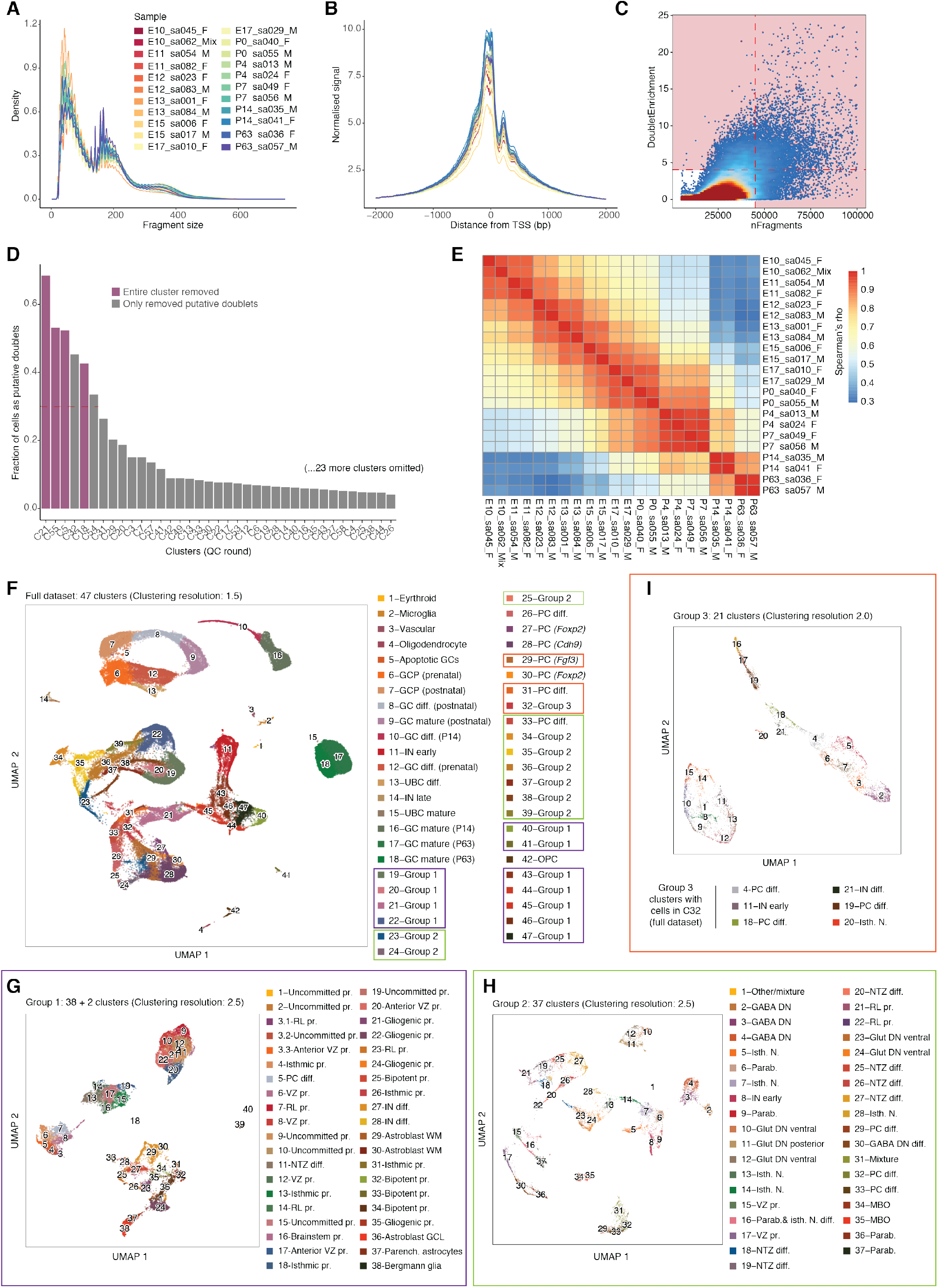
Quality control and processing of snATAC-seq data. (A) Fragment size distribution of the snATAC-seq libraries. (B) TSS enrichment scores of the snATAC-seq libraries. (C) 2D-density plot showing the number of fragments and doublet enrichment score per cell after 1^st^ pass removal of putative doublets. Cells in red-shaded areas were removed in the 2^nd^ round of doublet filtering. (D) Fraction of cells marked as doublets in the 2^nd^ filtering round per cluster. Clusters with more than 30% of doublets (horizontal line) were examined further for marker genes. Clusters marked in violet were removed completely. For all other clusters (gray) only putative doublets were removed. (E) Rank-based (Spearman’s rho) correlation coefficient of autosomal CRE accessibility across samples. (F) UMAP projection of 91,922 high quality cells colored by cluster assignment. Clusters that could not be resolved were combined in three groups (labeled as Group 1-3). Some annotated clusters (i.e., clusters 29, 31 and 33) were included in these groups to aid the split and annotation of the mixed clusters. The colored rectangles indicate which clusters were used for subclustering within each group. (G, H, I) UMAP projections of subclustering analyses for astroglia (Group 1; G), early-born neurons (Group 2; H) and GABAergic neurons at E13 (Group 3; I). Cell type abbreviations as in text; diff., differentiating; pr., progenitor.

**Fig. S2.**
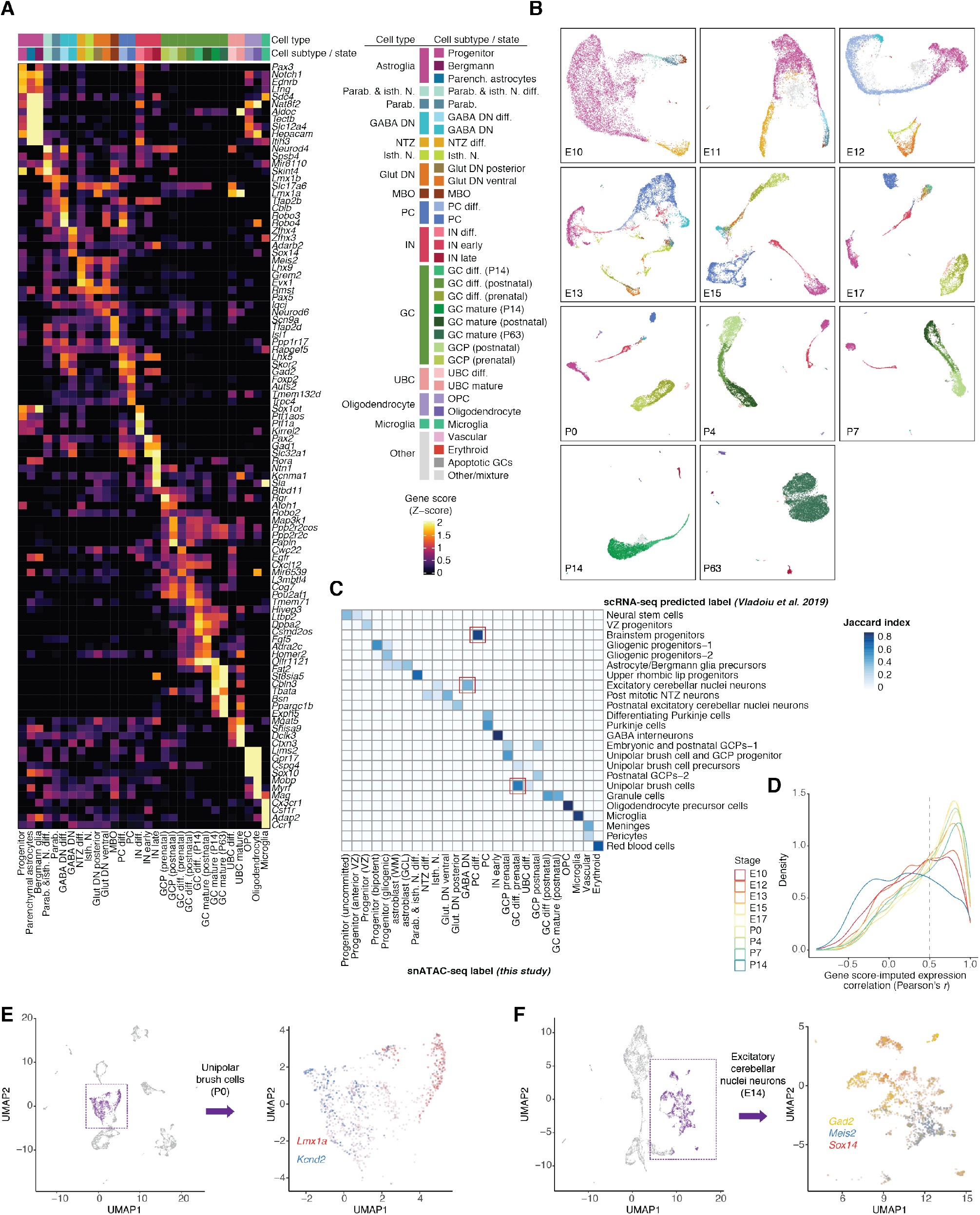
Cell type annotation and integration with scRNA-seq. (A) Activity scores of top 4 marker genes per cell type or state (Z-score, capped to 0-2). Marker genes were identified using Wilcoxon tests, as implemented in ArchR, and ranked based on the product of their log_2_ fold-change estimate and the –log_10_ *P*-value of the test. (B) UMAP projections and cell subtype/state annotations per developmental stage. (C) Jaccard similarity index between cell type labels from this study (columns) and transferred annotations after direct integration with scRNA-seq data (rows). Only labels with a similarity index of at least 0.15 with at least one other group are shown. The red rectangles mark unexpected matches (see Methods and E, F). (D) Per gene correlation (Pearson’s *r*) of mean gene score and imputed expression across snATAC-seq clusters for highly variable genes in the scRNA-seq data. The vertical line indicates the median correlation coefficient across developmental stages. (E) UMAP projection of 4,809 cells from P0 cerebellum profiled with scRNA-seq (*12*) (left). Cells originally annotated as UBCs are marked in purple and further examined for the expression of UBC (red) and GC (blue) specific markers (right). (F) UMAP projection of 6,068 cells from E14 cerebellum profiled with scRNA-seq (*12*) (left). Cells originally annotated as excitatory cerebellar nuclei neurons are marked in purple and further examined for the expression of *Meis2* (blue; marker of Glut. DN), *Gad2* (yellow; marker of GABAergic neurons) and *Sox14* (red; marker of GABA DN) (right).

**Fig. S3.**
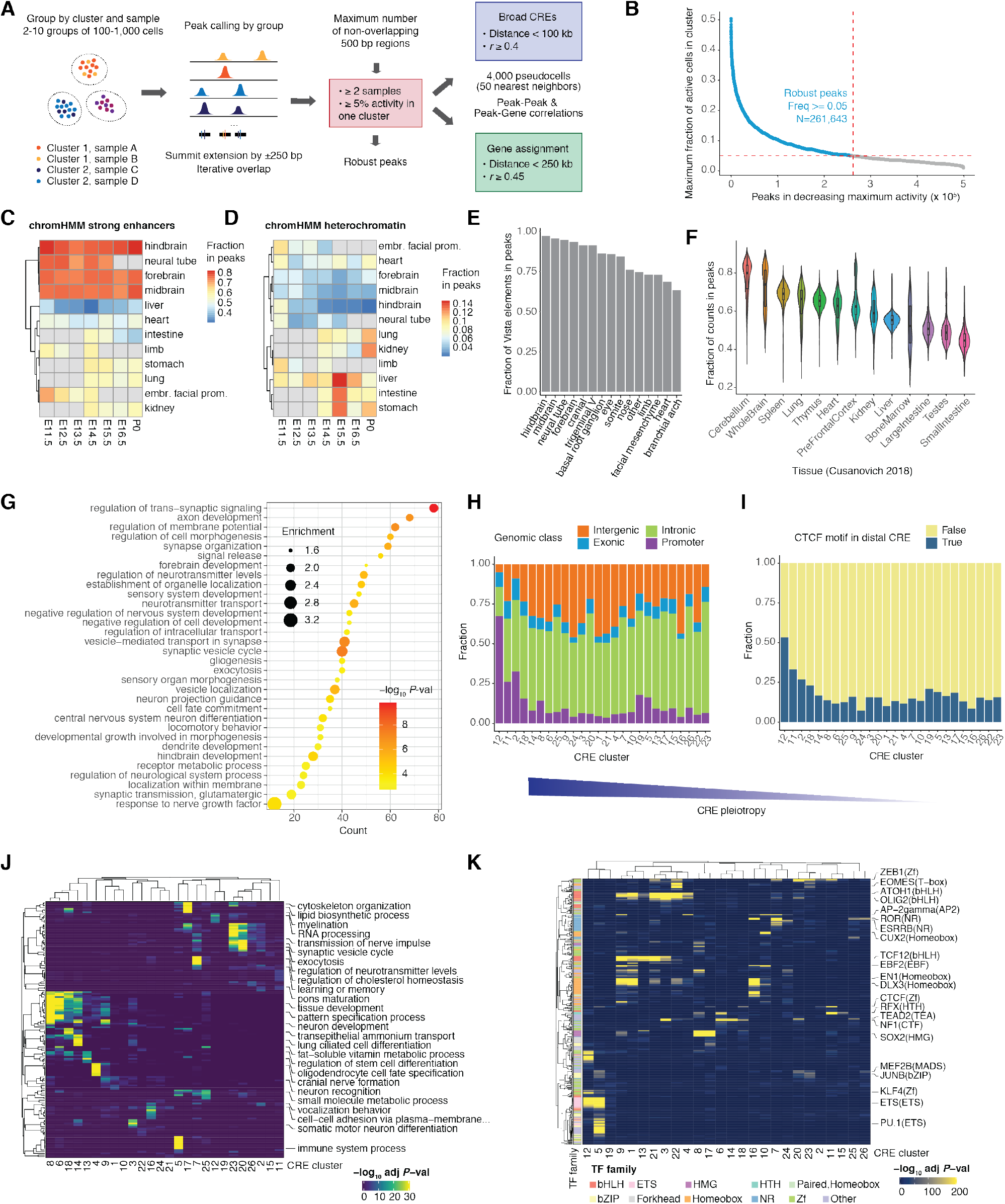
CRE identification, annotation and characterization. (A) Schematic representation of the procedure followed for the identification, filtering and assignment of CREs to their putative target genes and broader CRE units. (B) Putative CREs ranked based on their maximum activity across clusters (fraction of active cells in that cluster). Robust CREs were identified as active in at least 5% of the cells of at least one cluster. (C, D) Fraction of chromHMM predicted strong enhancers (C) and heterochromatin (D) across a series of tissues and developmental stages (*50*) overlapping robust CREs from this study. (E) Fraction of experimentally validated enhancers in mouse embryonic tissues (*49*) overlapping robust CREs from this study. Tissues are ordered by decreasing fraction. (F) Fraction of counts for cells from different adult mouse organs (*38*) in regions overlapping robust CREs from this study. Tissues are ordered by decreasing median fraction. (G) Biological process enrichment for genes associated with 5 or more non-redundant distal CREs. The x-axis indicates the number of genes associated with each term, the size and color of the dots the effect and significance of the enrichment based on a hypergeometric test. (H, I) Fraction of genomic classes (H) and distal CREs overlapping at least one CTCF motif (I) across clusters of CREs, indexed and ordered as in Fig. 2D. (J, K) Enrichment of biological processes of adjacent genes (J) and TF motifs (K) across clusters of CREs, indexed as in Fig. 2D (BH adjusted *P* < 0.05; hypergeometric test).

**Fig. S4.**
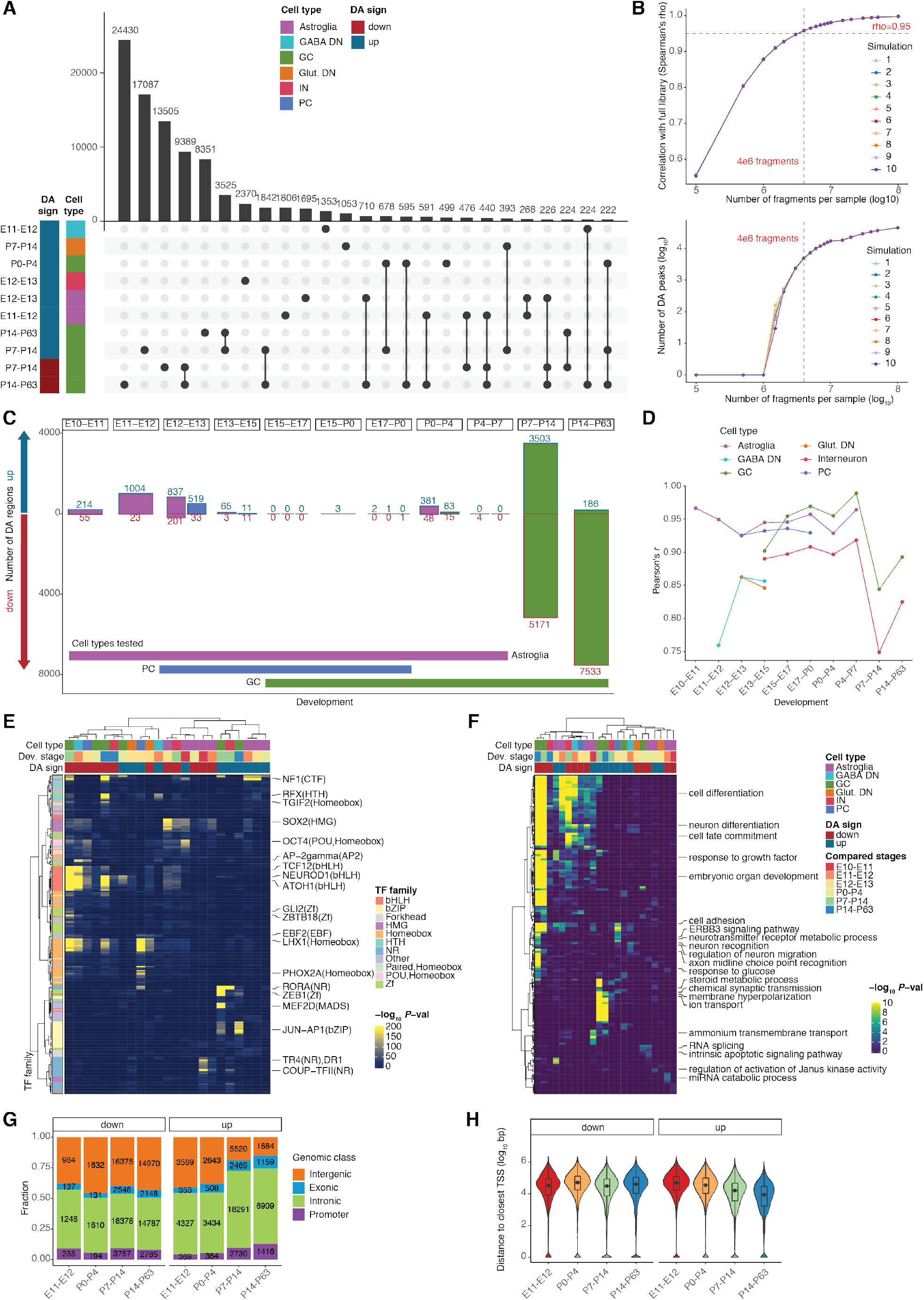
Identifying and characterizing periods of greater regulatory change. (A) Intersections across differentially accessible CREs from different cell types and developmental comparisons (left). Connected dots mark overlapping sets. The 10 largest sets (left) and 25 largest intersections (top) are shown. (B) Downsampling analysis of GCs in P14-P63. Spearman’s rho correlation with the full pseudobulk (mean value across the four samples) (top) and number of differentially accessible CREs detected between P14 and P63 (bottom). The vertical line indicates the number of fragments used for the analysis presented in (C). (C) Number of differentially accessible CREs across adjacent developmental stages after downsampling to the same number of fragments (4,000,000). Samples with fewer than 4,000,000 fragments were excluded. (D) Pearson’s correlation coefficients of CRE accessibility profiles from different cell types across adjacent developmental stages. (E, F) Enrichment of TF motifs (E) and biological processes of adjacent genes (F) for differentially accessible CREs, from Fig. 3A (BH adjusted *P* < 0.05; hypergeometric test). The cell type, developmental stage and sign of change are shown above the heatmaps. (G, H) Genomic class (G) and distance to closest TSS (H) for CREs identified as differentially accessible using data from all cell types, as shown in Fig. 3C.

**Fig. S5.**
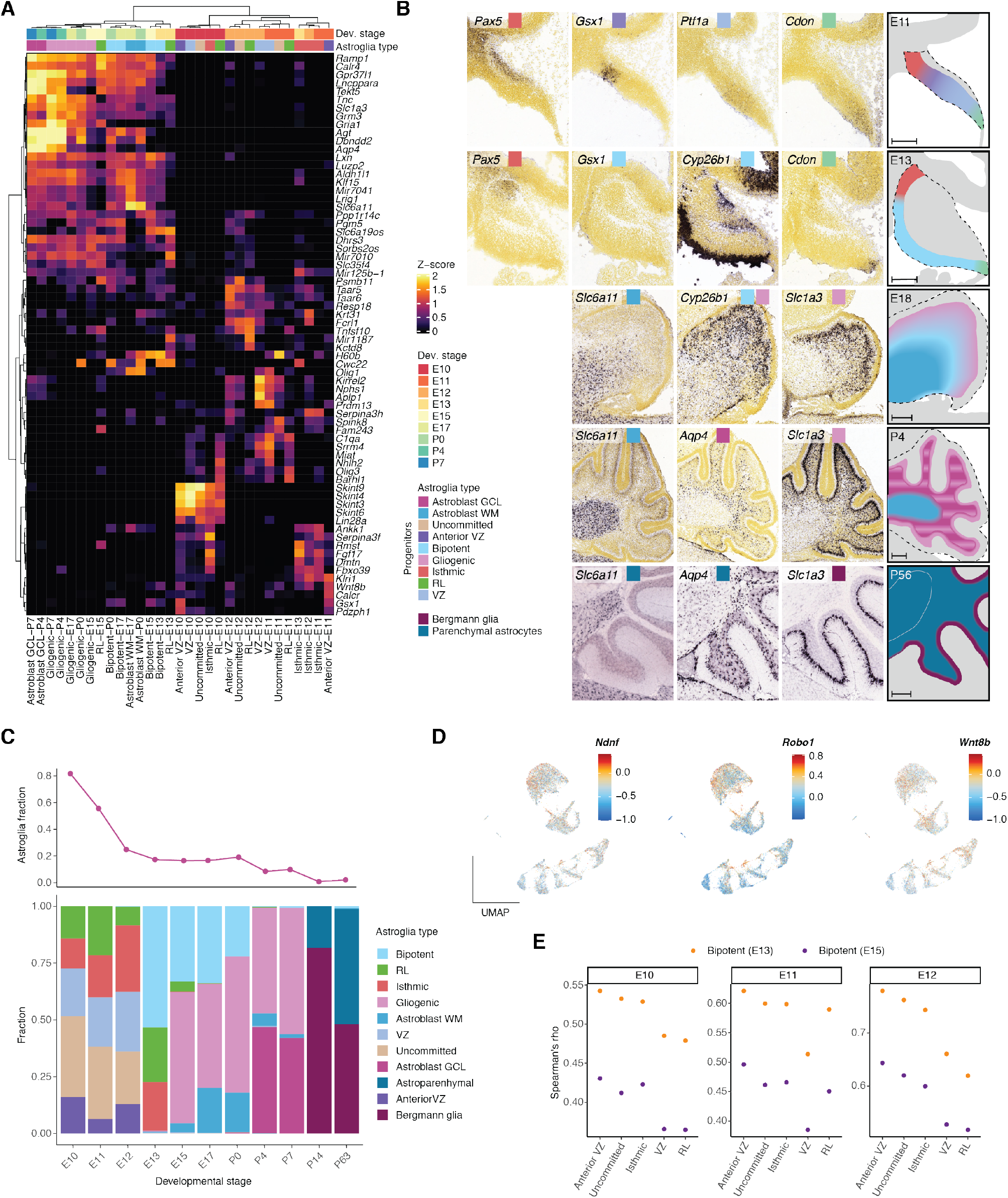
The regulatory landscape of progenitor cells in the developing cerebellum. (A) Activity scores of the top 4 marker genes per astroglia type or state (Z-score, capped to 0-2). Marker genes were identified using Wilcoxon tests as implemented in ArchR and ranked based on the product of their log_2_ fold-change estimate and the –log_10_ *P*-value of the test. (B) Estimated localization of astroglia types in the developing mouse cerebellum (right) based on *in situ* hybridization data (*58*) of selected marker genes (left). Sagittal sections counterstained with HP Yellow (E11-P4) or Nissl (P56) are shown. Scale bars: 200 μm. (C) Relative abundance of astroglia types (bottom) and overall fraction in the cerebellum (top) across developmental stages. (D) Gene activity scores for marker genes that are shared between anterior VZ and bipotent progenitors (Counts per 10^4^ fragments, capped at 10^th^ and 99^th^ quantiles and log_10_ transformed). (E) Spearman’s correlation coefficients based on global CRE activity between early (E10-E12) progenitor types and bipotent progenitors from E13 (orange) and E15 (purple).

**Fig. S6.**
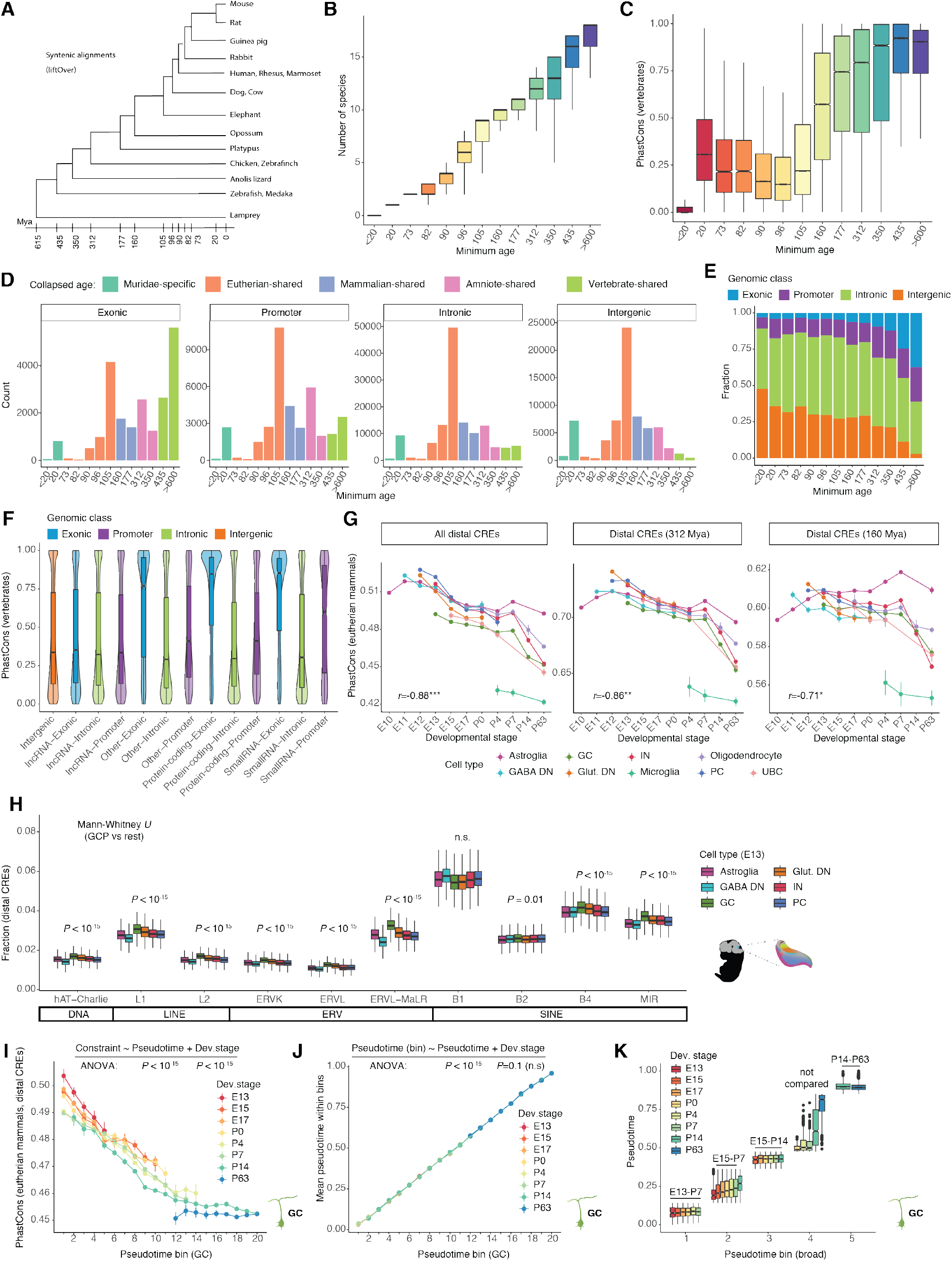
Evolutionary dynamics of CREs. (A) Species used in the syntenic alignments to infer the minimum age of CREs based on the date of divergence between mouse and the most distant species in which an alignment was detected. (B) Number of species in which a syntenic alignment was detected for CREs of different inferred ages. (C) Sequence constraint for CREs of different ages. (D) Number of CREs across genomic classes and age groups. Colors indicate broader age groups as used in Fig. 5E. (E) Relative representation of genomic classes across CRE age groups. (F,) Sequence constraint for CREs of different genomic classes. (G) Average sequence constraint across eutherian (commonly referred to as placental) mammals for all distal CREs (left), and subsets that originated 312 (middle) and 160 (right) million years ago (Mya), active across cell types and developmental stages. Vertical bars illustrate 95% confidence intervals of the estimates. Pearson’s *r* correlation coefficients between the estimates and development are shown (median across cell types; *P*<0.01**, *P*<0.001***). (H) Fraction of fragments in distal CREs overlapping transposable elements of different classes in cerebellar cell types at E13. (I, J) Average sequence constraint of distal CREs across eutherian mammals (J) and pseudotime value (K) along GC differentiation. Cells are separated by developmental stage and grouped in 20 pseudotime intervals with a step of 0.05. Vertical bars illustrate 95% confidence intervals of the estimates. (K) Distribution of pseudotime values across developmental stages and broad pseudotime bins (from Fig. 7B). Horizontal bars indicate the developmental stages that were used for temporal comparisons in matched differentiation states.

**Fig. S7.**
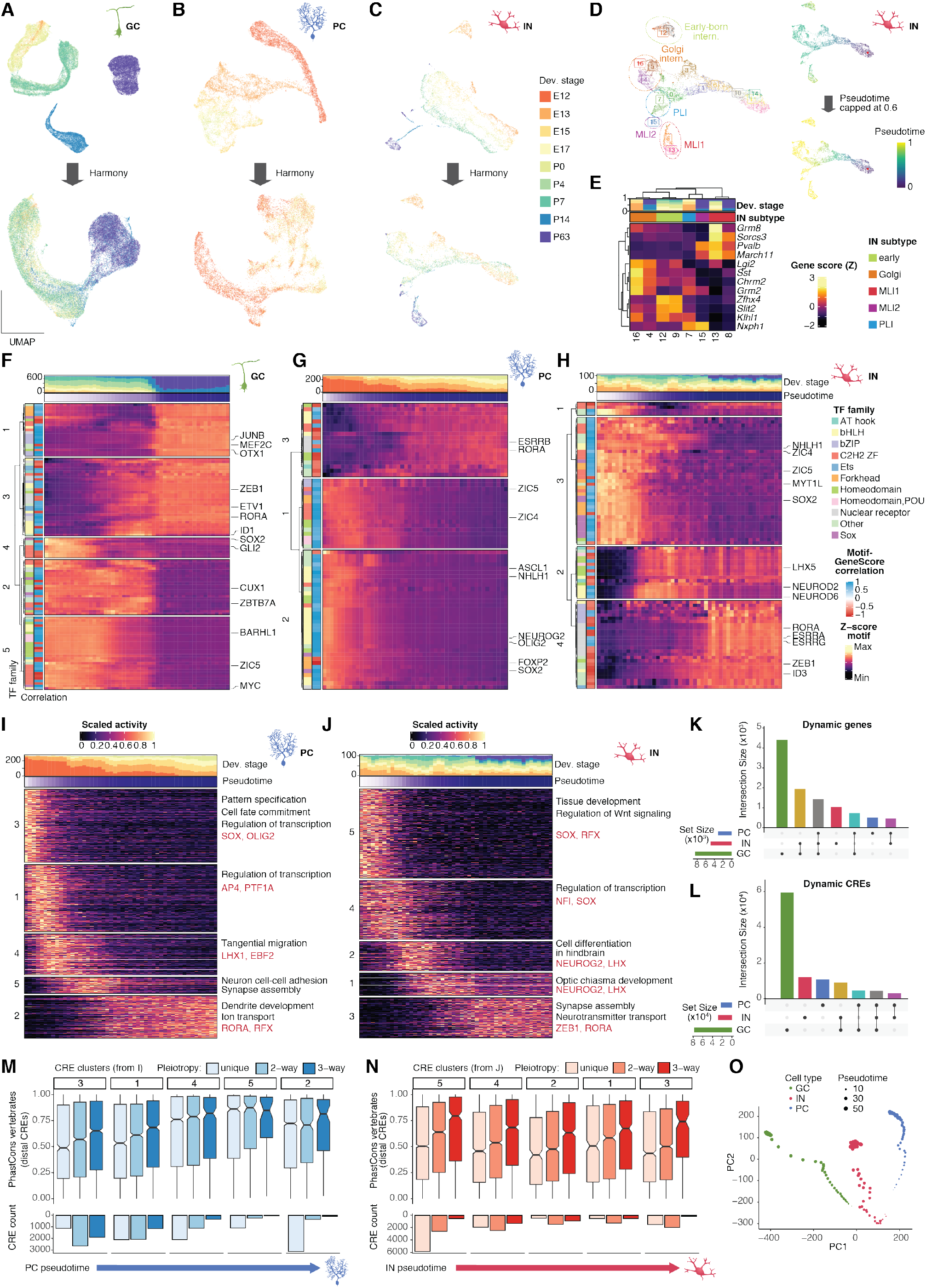
Chromatin accessibility during neuron differentiation. (A, B, C) UMAP projections of 35,153 GCs (A), 13,214 PCs (B) and 5,113 INs (C) before (top) and after (bottom) Harmony-alignment across developmental stages. Cells are colored by developmental stage. (D) UMAP projections of 5,113 Harmony-aligned INs colored by cluster (left), and pseudotime (right) before (top) and after (bottom) capping at 0.6 to eliminate differences between temporally specified subtypes (circled in the left plot). (E) Gene score activity (Z-score) of marker genes for mature IN clusters (as in D). Subtype annotation and relative contribution of developmental stages per cluster are shown above the heatmap. (F, G, H) Heatmap of Z-score scaled motif accessibility for TFs with dynamic activity across GC (F), PC (G) and IN (H) differentiation. Cells are grouped in 50 bins based on pseudotime ranks. The contribution of different developmental stages and the mean pseudotime value for each bin are shown above the heatmap. Motifs are grouped in clusters of similar activity. Information on the TF family and Pearson’s correlation between motif accessibility and gene score are shown (left). TFs with positive or negative correlations are classified as putative activators (blue) and repressors (red) respectively. (I, J) Heatmap of Z-score scaled activity of dynamic CREs across PC (I) and IN (J) differentiation. Cells are grouped in 50 bins based on pseudotime ranks. The contribution of different developmental stages and the mean pseudotime value for each bin are shown above the heatmap. Cluster numbers are indicated on the left. (K, L) Upset plots of intersections between genes (K) and CREs (L) with dynamic activity in GC, PC and IN differentiation. Connected dots mark overlapping sets. (M, N) Differences in sequence conservation (top) and abundance (bottom) of distal CREs with increasing pleiotropy (shading) active in different stages of PC (M) and IN (N) differentiation (indicated by cluster numbers as in I-J on top). (O) PCA of the chromatin accessibility landscape during GC (green), PC (blue) and IN (red) differentiation. For each neuron type, cells are grouped in 50 bins based on pseudotime ranks. Point size increases as pseudotime progresses.

## Supplementary Table legends

Table S1. Information on sample dissections, library preparation and statistics.

Table S2. Clustering, cell type annotation and summary statistics of high quality cells.

Table S3. TFs associated with the differentiation of major cerebellar neuron types.

Table S4. Biological process enrichments for CREs with differential accessibility across developmental stages in matched states of GC differentiation.

Table S5. Software and external resources used in this study.

## References

1. A. Sathyanesan, J. Zhou, J. Scafidi, D. H. Heck, R. V. Sillitoe, V. Gallo, Emerging connections between cerebellar development, behaviour and complex brain disorders. Nat. Rev. Neurosci. 20, 298–313 (2019).

2. K. Leto, M. Arancillo, E. B. E. Becker, A. Buffo, C. Chiang, B. Ding, W. B. Dobyns, I. Dusart, P. Haldipur, M. E. Hatten, M. Hoshino, A. L. Joyner, M. Kano, D. L. Kilpatrick, N. Koibuchi, S. Marino, S. Martinez, K. J. Millen, T. O. Millner, T. Miyata, E. Parmigiani, K. Schilling, G. Sekerková, R. V. Sillitoe, C. Sotelo, N. Uesaka, A. Wefers, R. J. T. Wingate, R. Hawkes, Consensus Paper: Cerebellar Development. Cerebellum. 15, 789–828 (2016).

3. J. J. White, R. V Sillitoe, Development of the cerebellum: From gene expression patterns to circuit maps. Wiley Interdiscip. Rev. Dev. Biol. 2, 149–164 (2013).

4. T. Butts, M. J. Green, R. J. T. Wingate, Development of the cerebellum: Simple steps to make a ‘little brain.’ Dev. 141 (2014), pp. 4031–4041.

5. J. Kebschull, N. Ringach, E. Richman, D. Friedmann, S. S. Kolluru, R. Jones, W. Allen, Y. Wang, H. Zhou, S. W. Cho, H. Chang, K. Deisseroth, S. Quake, L. Luo, Cerebellar nuclei evolved by repeatedly duplicating a conserved cell type set. Science (80-.). 370 (2020), doi:10.1126/science.abd5059.

6. T. Butts, M. S. Modrell, C. V. H. Baker, R. J. T. Wingate, The evolution of the vertebrate cerebellum: Absence of a proliferative external granule layer in a non-teleost ray-finned fish. Evol. Dev. 16, 92–100 (2014).

7. A. Iulianella, R. J. Wingate, C. B. Moens, E. Capaldo, The generation of granule cells during the development and evolution of the cerebellum. Dev. Dyn. 248, 506–513 (2019).

8. V. Y. Wang, H. Y. Zoghbi, Genetic regulation of cerebellar development. Nat. Rev. Neurosci. 2, 484–491 (2001).

9. M. Yamada, Y. Seto, S. Taya, T. Owa, Y. U. Inoue, T. Inoue, Y. Kawaguchi, Y. I. Nabeshima, M. Hoshino, Specification of spatial identities of cerebellar neuron progenitors by ptf1a and atoh1 for proper production of GABAergic and glutamatergic neurons. J. Neurosci. 34, 4786–4800 (2014).

10. Y. Seto, T. Nakatani, N. Masuyama, S. Taya, M. Kumai, Y. Minaki, A. Hamaguchi, Y. U. Inoue, T. Inoue, S. Miyashita, T. Fujiyama, M. Yamada, H. Chapman, K. Campbell, M. A. Magnuson, C. V. Wright, Y. Kawaguchi, K. Ikenaka, H. Takebayashi, S. Ishiwata, Y. Ono, M. Hoshino, Temporal identity transition from Purkinje cell progenitors to GABAergic interneuron progenitors in the cerebellum. Nat. Commun. 5 (2014), doi:10.1038/ncomms4337.

11. R. A. Carter, L. Bihannic, C. Rosencrance, J. L. Hadley, Y. Tong, T. N. Phoenix, S. Natarajan, J. Easton, P. A. Northcott, C. Gawad, A Single-Cell Transcriptional Atlas of the Developing Murine Cerebellum. Curr. Biol. 28, 2910–2920.e2 (2018).

12. M. C. Vladoiu, I. El-hamamy, L. K. Donovan, H. Farooq, B. L. Holgado, Y. Sundaravadanam, V. Ramaswamy, L. D. Hendrikse, S. Kumar, S. C. Mack, J. J. Y. Lee, V. Fong, K. Juraschka, D. Przelicki, A. Michealraj, P. Skowron, B. Luu, H. Suzuki, A. S. Morrissy, F. M. G. Cavalli, L. Garzia, C. Daniels, X. Wu, Childhood cerebellar tumours mirror conserved fetal transcriptional programs. Nature (2019), doi:10.1038/s41586-019-1158-7.

13. J. W. Wizeman, Q. Guo, E. M. Wilion, J. Y. H. Li, Specification of diverse cell types during early neurogenesis of the mouse cerebellum. Elife. 8, 1–24 (2019).

14. J. Peng, A. Sheng, Q. Xiao, L. Shen, X.-C. Ju, M. Zhang, S.-T. He, C. Wu, Z.-G. Luo, Single-cell transcriptomes reveal molecular specializations of neuronal cell types in the developing cerebellum. J. Mol. Cell Biol., 1–13 (2019).

15. K. A. Aldinger, Z. Thomson, P. Haldipur, M. Deng, A. E. Timms, M. Hirano, G. Santpere, C. Roco, A. B. Rosenberg, B. Lorente-Galdos, F. O. Gulden, D. O’Day, L. M. Overman, S. Lisgo, P. Alexandre, N. Sestan, D. Doherty, W. B. Dobyns, G. Seelig, I. A. Glass, K. J. Millen, bioRxiv, in press, doi:10.1101/2020.06.30.174391.

16. A. S. Nord, A. E. West, Neurobiological functions of transcriptional enhancers. Nat. Neurosci. 23 (2020), pp. 5–14.

17. H. K. Long, S. L. Prescott, J. Wysocka, Ever-Changing Landscapes: Transcriptional Enhancers in Development and Evolution. Cell. 167, 1170–1187 (2016).

18. S. Heinz, C. E. Romanoski, C. Benner, C. K. Glass, The selection and function of cell type-specific enhancers. Nat. Rev. Mol. Cell Biol. 16, 144–154 (2015).

19. D. Arendt, J. M. Musser, C. V. H. Baker, A. Bergman, C. Cepko, D. H. Erwin, M. Pavlicev, G. Schlosser, S. Widder, M. D. Laubichler, G. P. Wagner, The origin and evolution of cell types. Nat. Rev. Genet. 17, 744–757 (2016).

20. A. B. Stergachis, S. Neph, A. Reynolds, R. Humbert, B. Miller, S. L. Paige, B. Vernot, J. B. Cheng, R. E. Thurman, R. Sandstrom, E. Haugen, S. Heimfeld, C. E. Murry, J. M. Akey, J. A. Stamatoyannopoulos, Developmental fate and cellular maturity encoded in human regulatory DNA landscapes. Cell. 154, 888–903 (2013).

21. A. S. Nord, M. J. Blow, C. Attanasio, J. A. Akiyama, A. Holt, R. Hosseini, S. Phouanenavong, I. Plajzer-Frick, M. Shoukry, V. Afzal, J. L. R. Rubenstein, E. M. Rubin, L. A. Pennacchio, A. Visel, Rapid and pervasive changes in genome-wide enhancer usage during mammalian development. Cell. 155, 1521–1531 (2013).

22. D. U. Gorkin, I. Barozzi, Y. Zhao, Y. Zhang, H. Huang, A. Y. Lee, B. Li, J. Chiou, A. Wildberg, B. Ding, B. Zhang, M. Wang, J. S. Strattan, J. M. Davidson, Y. Qiu, V. Afzal, J. A. Akiyama, I. Plajzer-Frick, C. S. Novak, M. Kato, T. H. Garvin, Q. T. Pham, A. N. Harrington, B. J. Mannion, E. A. Lee, Y. Fukuda-Yuzawa, Y. He, S. Preissl, S. Chee, J. Y. Han, B. A. Williams, D. Trout, H. Amrhein, H. Yang, J. M. Cherry, W. Wang, K. Gaulton, J. R. Ecker, Y. Shen, D. E. Dickel, A. Visel, L. A. Pennacchio, B. Ren, An atlas of dynamic chromatin landscapes in mouse fetal development. Nature. 583, 744–751 (2020).

23. Y. He, M. Hariharan, D. U. Gorkin, D. E. Dickel, C. Luo, R. G. Castanon, J. R. Nery, A. Y. Lee, Y. Zhao, H. Huang, B. A. Williams, D. Trout, H. Amrhein, R. Fang, H. Chen, B. Li, A. Visel, L. A. Pennacchio, B. Ren, J. R. Ecker, Spatiotemporal DNA methylome dynamics of the developing mouse fetus. Nature. 583, 752–759 (2020).

24. C. L. Frank, F. Liu, R. Wijayatunge, L. Song, M. T. Biegler, M. G. Yang, C. M. Vockley, A. Safi, C. A. Gersbach, G. E. Crawford, A. E. West, Regulation of chromatin accessibility and Zic binding at enhancers in the developing cerebellum. Nat. Neurosci. 18, 647–656 (2015).

25. D. Villar, C. Berthelot, S. Aldridge, T. F. Rayner, M. Lukk, M. Pignatelli, T. J. Park, R. Deaville, J. T. Erichsen, A. J. Jasinska, J. M. A. Turner, M. F. Bertelsen, E. P. Murchison, P. Flicek, D. T. Odom, Enhancer evolution across 20 mammalian species. Cell. 160, 554–566 (2015).

26. C. Berthelot, D. Villar, J. E. Horvath, D. T. Odom, P. Flicek, Complexity and conservation of regulatory landscapes underlie evolutionary resilience of mammalian gene expression. Nat. Ecol. Evol. 2, 152–163 (2018).

27. J. Cotney, J. Leng, J. Yin, S. K. Reilly, L. E. Demare, D. Emera, A. E. Ayoub, P. Rakic, J. P. Noonan, The evolution of lineage-specific regulatory activities in the human embryonic limb. Cell. 154, 185 (2013).

28. S. K. Reilly, J. Yin, A. E. Ayoub, D. Emera, J. Leng, J. Cotney, R. Sarro, P. Rakic, J. P. Noonan, Evolutionary changes in promoter and enhancer activity during human corticogenesis. Sci. (New York, NY). 347, 1155–1159 (2015).

29. D. Brawand, M. Soumillon, A. Necsulea, P. Julien, G. Csárdi, P. Harrigan, M. Weier, A. Liechti, A. Aximu-Petri, M. Kircher, F. W. Albert, U. Zeller, P. Khaitovich, F. Grützner, S. Bergmann, R. Nielsen, S. Pääbo, H. Kaessmann, The evolution of gene expression levels in mammalian organs. Nature. 478, 343–348 (2011).

30. M. Cardoso-Moreira, J. Halbert, D. Valloton, B. Velten, C. Chen, Y. Shao, A. Liechti, K. Ascenção, C. Rummel, S. Ovchinnikova, P. V. Mazin, I. Xenarios, K. Harshman, M. Mort, D. N. Cooper, C. Sandi, M. J. Soares, P. G. Ferreira, S. Afonso, M. Carneiro, J. M. A. Turner, J. L. VandeBerg, A. Fallahshahroudi, P. Jensen, R. Behr, S. Lisgo, S. Lindsay, P. Khaitovich, W. Huber, J. Baker, S. Anders, Y. E. Zhang, H. Kaessmann, Gene expression across mammalian organ development. Nature. 571, 505–509 (2019).

31. G. Bejerano, M. Pheasant, I. Makunin, S. Stephen, W. J. Kent, J. S. Mattick, D. Haussler, Ultraconserved elements in the human genome. Science (80-.). 304, 1321–1325 (2004).

32. D. E. Dickel, A. R. Ypsilanti, R. Pla, Y. Zhu, I. Barozzi, B. J. Mannion, Y. S. Khin, Y. Fukuda-Yuzawa, I. Plajzer-Frick, C. S. Pickle, E. A. Lee, A. N. Harrington, Q. T. Pham, T. H. Garvin, M. Kato, M. Osterwalder, J. A. Akiyama, V. Afzal, J. L. R. Rubenstein, L. A. Pennacchio, A. Visel, Ultraconserved Enhancers Are Required for Normal Development. Cell. 172, 491–499.e15 (2018).

33. I. Sarropoulos, R. Marin, M. Cardoso-Moreira, H. Kaessmann, Developmental dynamics of lncRNAs across mammalian organs and species. Nature. 571, 510–514 (2019).

34. J. D. Buenrostro, P. G. Giresi, L. C. Zaba, H. Y. Chang, W. J. Greenleaf, Transposition of native chromatin for fast and sensitive epigenomic profiling of open chromatin, DNA-binding proteins and nucleosome position. Nat. Methods. 10, 1213–1218 (2013).

35. J. D. Buenrostro, B. Wu, U. M. Litzenburger, D. Ruff, M. L. Gonzales, M. P. Snyder, H. Y. Chang, W. J. Greenleaf, Single-cell chromatin accessibility reveals principles of regulatory variation. Nature. 523, 486–490 (2015).

36. A. T. Satpathy, J. M. Granja, K. E. Yost, Y. Qi, F. Meschi, G. P. McDermott, B. N. Olsen, M. R. Mumbach, S. E. Pierce, M. R. Corces, P. Shah, J. C. Bell, D. Jhutty, C. M. Nemec, J. Wang, L. Wang, Y. Yin, P. G. Giresi, A. L. S. Chang, G. X. Y. Zheng, W. J. Greenleaf, H. Y. Chang, Massively parallel single-cell chromatin landscapes of human immune cell development and intratumoral T cell exhaustion. Nat. Biotechnol. 37, 925–936 (2019).

37. C. A. Lareau, F. M. Duarte, J. G. Chew, V. K. Kartha, Z. D. Burkett, A. S. Kohlway, D. Pokholok, M. J. Aryee, F. J. Steemers, R. Lebofsky, J. D. Buenrostro, Droplet-based combinatorial indexing for massive-scale single-cell chromatin accessibility. Nat. Biotechnol. 37, 916–924 (2019).

38. D. A. Cusanovich, A. J. Hill, D. Aghamirzaie, R. M. Daza, H. A. Pliner, J. B. Berletch, G. N. Filippova, X. Huang, L. Christiansen, W. S. DeWitt, C. Lee, S. G. Regalado, D. F. Read, F. J. Steemers, C. M. Disteche, C. Trapnell, J. Shendure, A Single-Cell Atlas of In Vivo Mammalian Chromatin Accessibility. Cell, 1–16 (2018).

39. Y. E. Li, S. Preissl, X. Hou, Z. Zhang, K. Zhang, R. Fang, Y. Qiu, O. Poirion, B. Li, H. Liu, X. Wang, J. Y. Han, J. Lucero, Y. Yan, S. Kuan, D. Gorkin, M. Nunn, E. A. Mukamel, M. M. Behrens, J. Ecker, B. Ren, bioRxiv, in press, doi:10.1101/2020.05.10.087585.

40. S. Preissl, R. Fang, H. Huang, Y. Zhao, R. Raviram, D. U. Gorkin, Y. Zhang, B. C. Sos, V. Afzal, D. E. Dickel, S. Kuan, A. Visel, L. A. Pennacchio, K. Zhang, B. Ren, Single-nucleus analysis of accessible chromatin in developing mouse forebrain reveals cell-type-specific transcriptional regulation. Nat. Neurosci. (2018), doi:10.1038/s41593-018-0079-3.

41. D. Di Bella, E. Habibi, S. Yang, R. Stickels, J. Brown, P. Yadollahpour, F. Chen, E. Macosko, A. Regev, P. Arlotta, bioRxiv, in press, doi:10.1101/2020.07.02.185439.

42. R. S. Ziffra, C. N. Kim, A. Wilfert, T. N. Turner, M. Haeussler, A. M. Casella, P. F. Przytycki, A. Kreimer, K. S. Pollard, S. A. Ament, E. E. Eichler, N. Ahituv, T. J. Nowakowski, bioRxiv, in press, doi:10.1101/2019.12.30.891549.

43. S. Domcke, A. J. Hill, R. M. Daza, J. Cao, D. R. O’Day, H. A. Pliner, K. A. Aldinger, D. Pokholok, F. Zhang, J. H. Milbank, M. A. Zager, I. A. Glass, F. J. Steemers, D. Doherty, C. Trapnell, D. A. Cusanovich, J. Shendure, A human cell atlas of fetal chromatin accessibility. Science. 370 (2020), doi:10.1126/science.aba7612.

44. J. Granja, M. R. Corces, S. Pierce, S. T. Bagdatli, H. Choudhry, H. Chang, W. Greenleaf, ArchR: An integrative and scalable software package for single-cell chromatin accessibility analysis. bioRxiv (2020), doi:10.1101/2020.04.28.066498.

45. K. J. Millen, E. Y. Steshina, I. Y. Iskusnykh, V. V. Chizhikov, Transformation of the cerebellum into more ventral brainstem fates causes cerebellar agenesis in the absence of Ptf1a function. Proc. Natl. Acad. Sci. U. S. A. 111 (2014), doi:10.1073/pnas.1315024111.

46. R. Machold, G. Fishell, Math1 is expressed in temporally discrete pools of cerebellar rhombic-lip neural progenitors. Neuron. 48, 17–24 (2005).

47. A. Kriegstein, A. Alvarez-Buylla, The glial nature of embryonic and adult neural stem cells. Annu. Rev. Neurosci. 32, 149–184 (2009).

48. F. Doetsch, The glial identity of neural stem cells. Nat. Neurosci. 6, 1127–1134 (2003).

49. A. Visel, S. Minovitsky, I. Dubchak, L. A. Pennacchio, VISTA Enhancer Browser - A database of tissue-specific human enhancers. Nucleic Acids Res. 35, D88–D92 (2007).

50. The ENCODE Project Consortium, Expanded encyclopaedias of DNA elements in the human and mouse genomes. Nature. 583, 699–710 (2020).

51. G. Sabarís, I. Laiker, E. Preger-Ben Noon, N. Frankel, Actors with Multiple Roles: Pleiotropic Enhancers and the Paradigm of Enhancer Modularity. Trends Genet. 35, 423–433 (2019).

52. W. Meuleman, A. Muratov, E. Rynes, J. Halow, K. Lee, D. Bates, M. Diegel, D. Dunn, F. Neri, A. Teodosiadis, A. Reynolds, E. Haugen, J. Nelson, A. Johnson, M. Frerker, M. Buckley, R. Sandstrom, J. Vierstra, R. Kaul, J. Stamatoyannopoulos, Index and biological spectrum of human DNase I hypersensitive sites. Nature. 584 (2020), doi:10.1038/s41586-020-2559-3.

53. M. J. van Essen, S. Nayler, E. B. E. Becker, J. Jacob, Deconstructing cerebellar development cell by cell. PLoS Genet. 16 (2020), p. e1008630.

54. T. Zhang, T. Liu, N. Mora, J. Guegan, M. Bertrand, X. Contreras, A. H. Hansen, C. Streicher, M. Anderle, L. Tiberi, S. Hippenmeyer, B. A. Hassan, bioRxiv, in press, doi:10.1101/2020.03.18.997205.

55. E. Parmigiani, K. Leto, C. Rolando, M. Figueres-Oñate, L. López-Mascaraque, A. Buffo, F. Rossi, Heterogeneity and bipotency of astroglial-like cerebellar progenitors along the interneuron and glial lineages. J. Neurosci. 35, 7388–7402 (2015).

56. V. Cerrato, E. Parmigiani, M. Figueres-Oñate, M. Betizeau, J. Aprato, I. Nanavaty, P. Berchialla, F. Luzzati, C. de’Sperati, L. López-Mascaraque, A. Buffo, Multiple origins and modularity in the spatiotemporal emergence of cerebellar astrocyte heterogeneity (2018), vol. 16.

57. A. Sagner, I. Zhang, T. Watson, J. Lazaro, M. Melchionda, J. Briscoe, Temporal patterning of the central nervous system by a shared transcription factor code. bioRxiv (2020), doi:https://doi.org/10.1101/2020.11.10.376491.

58. Allen Institute for Brain Science, Allen Developing Mouse Brain Atlas (2008), (available at http://developingmouse.brain-map.org/).

59. D. E. Wagner, A. M. Klein, Lineage tracing meets single-cell omics: opportunities and challenges. Nat. Rev. Genet., 1–18 (2020).

60. A. Siepel, G. Bejerano, J. S. Pedersen, A. S. Hinrichs, M. Hou, K. Rosenbloom, H. Clawson, J. Spieth, L. D. W. Hillier, S. Richards, G. M. Weinstock, R. K. Wilson, R. A. Gibbs, W. J. Kent, W. Miller, D. Haussler, Evolutionarily conserved elements in vertebrate, insect, worm, and yeast genomes. Genome Res. 15, 1034–1050 (2005).

61. L. Geirsdottir, E. David, H. Keren-Shaul, A. Weiner, S. C. Bohlen, J. Neuber, A. Balic, A. Giladi, F. Sheban, C. A. Dutertre, C. Pfeifle, F. Peri, A. Raffo-Romero, J. Vizioli, K. Matiasek, C. Scheiwe, S. Meckel, K. Mätz-Rensing, F. van der Meer, F. R. Thormodsson, C. Stadelmann, N. Zilkha, T. Kimchi, F. Ginhoux, I. Ulitsky, D. Erny, I. Amit, M. Prinz, Cross-Species Single-Cell Analysis Reveals Divergence of the Primate Microglia Program. Cell. 179, 1609–1622.e16 (2019).

62. A. Necsulea, H. Kaessmann, Evolutionary dynamics of coding and non-coding transcriptomes. Nat. Rev. Genet. 15, 734–748 (2014).

63. A. F. A. Smit, A. D. Riggs, MIRs are classic, tRNA-derived SINEs that amplified before the mammalian radiation. Nucleic Acids Res. 23, 98–102 (1995).

64. L. Haghverdi, M. Büttner, F. A. Wolf, F. Buettner, F. J. Theis, Diffusion pseudotime robustly reconstructs lineage branching. Nat. Methods. 13, 845–848 (2016).

65. V. Kozareva, C. Martin, T. Osorno, S. Rudolph, C. Guo, C. Vanderburg, N. Nadaf, A. Regev, W. Regehr, E. Macosko, A transcriptomic atlas of the mouse cerebellum reveals regional specializations and novel cell types. bioRxiv (2020), doi:10.1101/2020.03.04.976407.

66. A. N. Schep, B. Wu, J. D. Buenrostro, W. J. Greenleaf, ChromVAR: Inferring transcription-factor-associated accessibility from single-cell epigenomic data. Nat. Methods. 14, 975–978 (2017).

67. I. Berest, C. Arnold, A. Reyes-Palomares, G. Palla, K. D. Rasmussen, H. Giles, P. M. Bruch, W. Huber, S. Dietrich, K. Helin, J. B. Zaugg, Quantification of Differential Transcription Factor Activity and Multiomics-Based Classification into Activators and Repressors: diffTF. Cell Rep. 29, 3147–3159.e12 (2019).

68. L. Telley, G. Agirman, J. Prados, N. Amberg, S. Fièvre, P. Oberst, G. Bartolini, I. Vitali, C. Cadilhac, S. Hippenmeyer, L. Nguyen, A. Dayer, D. Jabaudon, Temporal patterning of apical progenitors and their daughter neurons in the developing neocortex. Science (80-.). 364 (2019), doi:10.1126/science.aav2522.

69. R. Andersson, C. Gebhard, I. Miguel-Escalada, I. Hoof, J. Bornholdt, M. Boyd, Y. Chen, X. Zhao, C. Schmidl, T. Suzuki, E. Ntini, E. Arner, E. Valen, K. Li, L. Schwarzfischer, D. Glatz, J. Raithel, B. Lilje, N. Rapin, F. O. Bagger, M. Jørgensen, P. R. Andersen, N. Bertin, O. Rackham, A. M. Burroughs, J. K. Baillie, Y. Ishizu, Y. Shimizu, E. Furuhata, S. Maeda, Y. Negishi, C. J. Mungall, T. F. Meehan, T. Lassmann, M. Itoh, H. Kawaji, N. Kondo, J. Kawai, A. Lennartsson, C. O. Daub, P. Heutink, D. A. Hume, T. H. Jensen, H. Suzuki, Y. Hayashizaki, F. Müller, A. R. R. Forrest, P. Carninci, M. Rehli, A. Sandelin, An atlas of active enhancers across human cell types and tissues. Nature. 507, 455–461 (2014).

70. H. S. Rhee, M. Closser, Y. Guo, E. V. Bashkirova, G. C. Tan, D. K. Gifford, H. Wichterle, Expression of Terminal Effector Genes in Mammalian Neurons Is Maintained by a Dynamic Relay of Transient Enhancers. Neuron. 92, 1252–1265 (2016).

71. E. Legué, J. L. Gottshall, E. Jaumouillé, A. Roselló-Díez, W. Shi, L. H. Barraza, S. Washington, R. L. Grant, A. L. Joyner, Differential timing of granule cell production during cerebellum development underlies generation of the foliation pattern. Neural Dev. 11, 1–14 (2016).

72. J. S. Espinosa, L. Luo, Timing neurogenesis and differentiation: Insights from quantitative clonal analyses of cerebellar granule cells. J. Neurosci. 28, 2301–2312 (2008).

73. S. L. Prescott, R. Srinivasan, M. C. Marchetto, I. Grishina, I. Narvaiza, L. Selleri, F. H. Gage, T. Swigut, J. Wysocka, Enhancer Divergence and cis-Regulatory Evolution in the Human and Chimp Neural Crest. Cell. 163, 68–84 (2015).

74. N. Dukler, Y. F. Huang, A. Siepel, S. Kosakovsky Pond, Phylogenetic Modeling of Regulatory Element Turnover Based on Epigenomic Data. Mol. Biol. Evol. 37, 2137–2152 (2020).

75. A. T. Kalinka, K. M. Varga, D. T. Gerrard, S. Preibisch, D. L. Corcoran, J. Jarrells, U. Ohler, C. M. Bergman, P. Tomancak, Gene expression divergence recapitulates the developmental hourglass model. Nature. 468, 811–816 (2010).

76. A. Abzhanov, Von Baer’s law for the ages: Lost and found principles of developmental evolution. Trends Genet. 29 (2013), pp. 712–722.

77. M. Götz, S. Sirko, J. Beckers, M. Irmler, Reactive astrocytes as neural stem or progenitor cells: In vivo lineage, In vitro potential, and Genome-wide expression analysis. Glia. 63, 1452–1468 (2015).

78. A. Volterra, J. Meldolesi, Astrocytes, from brain glue to communication elements: The revolution continues. Nat. Rev. Neurosci. 6, 626–640 (2005).

79. S. R. Krishnaswami, R. V Grindberg, M. Novotny, P. Venepally, B. Lacar, K. Bhutani, S. B. Linker, S. Pham, J. A. Erwin, J. A. Miller, R. Hodge, J. K. McCarthy, M. Kelder, J. McCorrison, B. D. Aevermann, F. D. Fuertes, R. H. Scheuermann, J. Lee, E. S. Lein, N. Schork, M. J. McConnell, F. H. Gage, R. S. Lasken, Using single nuclei for RNA-seq to capture the transcriptome of postmortem neurons. Nat. Protoc. 11, 499–524 (2016).

80. D. A. Cusanovich, R. Daza, A. Adey, H. A. Pliner, L. Christiansen, K. L. Gunderson, F. J. Steemers, C. Trapnell, J. Shendure, Multiplex single-cell profiling of chromatin accessibility by combinatorial cellular indexing. Science (80-.). 348, 910–914 (2015).

81. T. Stuart, A. Butler, P. Hoffman, C. Hafemeister, E. Papalexi, W. M. Mauck, Y. Hao, M. Stoeckius, P. Smibert, R. Satija, Comprehensive Integration of Single-Cell Data. Cell. 177, 1888–1902.e21 (2019).

82. I. Korsunsky, N. Millard, J. Fan, K. Slowikowski, F. Zhang, K. Wei, Y. Baglaenko, M. Brenner, P. Loh, S. Raychaudhuri, Fast, sensitive and accurate integration of single-cell data with Harmony. Nat. Methods. 16, 1289–1296 (2019).

83. H. T. Prekop, A. Kroiss, V. Rook, L. Zagoraiou, T. M. Jessell, C. Fernandes, A. Delogu, R. J. T. Wingate, Sox14 is required for a specific subset of cerebello–olivary projections. J. Neurosci. 38, 9539–9550 (2018).

84. V. V. Chizhikov, A. G. Lindgren, Y. Mishima, R. W. Roberts, K. A. Aldinger, G. R. Miesegaes, D. Spencer Currle, E. S. Monuki, K. J. Millen, Lmx1a regulates fates and location of cells originating from the cerebellar rhombic lip and telencephalic cortical hem. Proc. Natl. Acad. Sci. U. S. A. 107, 10725–10730 (2010).

85. Y. Zhang, T. Liu, C. A. Meyer, J. Eeckhoute, D. S. Johnson, B. E. Bernstein, C. Nussbaum, R. M. Myers, M. Brown, W. Li, X. S. Shirley, Model-based analysis of ChIP-Seq (MACS). Genome Biol. 9, R137 (2008).

86. M. Haeussler, A. S. Zweig, C. Tyner, M. L. Speir, K. R. Rosenbloom, B. J. Raney, C. M. Lee, B. T. Lee, A. S. Hinrichs, J. N. Gonzalez, D. Gibson, M. Diekhans, H. Clawson, J. Casper, G. P. Barber, D. Haussler, R. M. Kuhn, W. J. Kent, The UCSC Genome Browser database: 2019 update. Nucleic Acids Res. 47, D853–D858 (2019).

87. F. Cunningham, P. Achuthan, W. Akanni, J. Allen, M. R. Amode, I. M. Armean, R. Bennett, J. Bhai, K. Billis, S. Boddu, C. Cummins, C. Davidson, K. J. Dodiya, A. Gall, C. G. Girón, L. Gil, T. Grego, L. Haggerty, E. Haskell, T. Hourlier, O. G. Izuogu, S. H. Janacek, T. Juettemann, M. Kay, M. R. Laird, I. Lavidas, Z. Liu, J. E. Loveland, J. C. Marugán, T. Maurel, A. C. McMahon, B. Moore, J. Morales, J. M. Mudge, M. Nuhn, D. Ogeh, A. Parker, A. Parton, M. Patricio, A. I. Abdul Salam, B. M. Schmitt, H. Schuilenburg, D. Sheppard, H. Sparrow, E. Stapleton, M. Szuba, K. Taylor, G. Threadgold, A. Thormann, A. Vullo, B. Walts, A. Winterbottom, A. Zadissa, M. Chakiachvili, A. Frankish, S. E. Hunt, M. Kostadima, N. Langridge, F. J. Martin, M. Muffato, E. Perry, M. Ruffier, D. M. Staines, S. J. Trevanion, B. L. Aken, A. D. Yates, D. R. Zerbino, P. Flicek, Ensembl 2019. Nucleic Acids Res. 47, D745–D751 (2019).

88. A. R. Quinlan, I. M. Hall, BEDTools: A flexible suite of utilities for comparing genomic features. Bioinformatics. 26, 841–842 (2010).

89. M. T. Weirauch, A. Yang, M. Albu, A. G. Cote, A. Montenegro-Montero, P. Drewe, H. S. Najafabadi, S. A. Lambert, I. Mann, K. Cook, H. Zheng, A. Goity, H. van Bakel, J. C. Lozano, M. Galli, M. G. Lewsey, E. Huang, T. Mukherjee, X. Chen, J. S. Reece-Hoyes, S. Govindarajan, G. Shaulsky, A. J. M. Walhout, F. Y. Bouget, G. Ratsch, L. F. Larrondo, J. R. Ecker, T. R. Hughes, Determination and inference of eukaryotic transcription factor sequence specificity. Cell. 158, 1431–1443 (2014).

90. W. J. Kent, A. S. Zweig, G. Barber, A. S. Hinrichs, D. Karolchik, BigWig and BigBed: Enabling browsing of large distributed datasets. Bioinformatics. 26, 2204–2207 (2010).

91. D. Karolchik, The UCSC Table Browser data retrieval tool. Nucleic Acids Res. 32, 493D – 496 (2004).

92. D. Hnisz, B. J. Abraham, T. I. Lee, A. Lau, V. Saint-André, A. A. Sigova, H. A. Hoke, R. A. Young, Super-enhancers in the control of cell identity and disease. Cell. 155, 934 (2013).

93. W. A. Whyte, D. A. Orlando, D. Hnisz, B. J. Abraham, C. Y. Lin, M. H. Kagey, P. B. Rahl, T. I. Lee, R. A. Young, Master transcription factors and mediator establish super-enhancers at key cell identity genes. Cell. 153, 307–319 (2013).

94. J. M. Granja, S. Klemm, L. M. McGinnis, A. S. Kathiria, A. Mezger, M. R. Corces, B. Parks, E. Gars, M. Liedtke, G. X. Y. Zheng, H. Y. Chang, R. Majeti, W. J. Greenleaf, Single-cell multiomic analysis identifies regulatory programs in mixed-phenotype acute leukemia. Nat. Biotechnol. 37 (2019), pp. 1458–1465.

95. C. Y. McLean, D. Bristor, M. Hiller, S. L. Clarke, B. T. Schaar, C. B. Lowe, A. M. Wenger, G. Bejerano, GREAT improves functional interpretation of cis-regulatory regions. Nat. Biotechnol. 28, 495–501 (2010).

96. S. Heinz, C. Benner, N. Spann, E. Bertolino, Y. C. Lin, P. Laslo, J. X. Cheng, C. Murre, H. Singh, C. K. Glass, Simple Combinations of Lineage-Determining Transcription Factors Prime cis-Regulatory Elements Required for Macrophage and B Cell Identities. Mol. Cell. 38, 576–589 (2010).

97. F. Supek, M. Bošnjak, N. Škunca, T. Šmuc, Revigo summarizes and visualizes long lists of gene ontology terms. PLoS One. 6 (2011), doi:10.1371/journal.pone.0021800.

98. A. E. Trevino, N. Sinnott-Armstrong, J. Andersen, S. Yoon, N. Huber, J. K. Pritchard, H. Y. Chang, W. J. Greenleaf, S. P. Paşca, Chromatin accessibility dynamics in a model of human forebrain development. Science. 367 (2020), doi:10.1126/science.aay1645.

99. M. I. Love, W. Huber, S. Anders, Moderated estimation of fold change and dispersion for RNA-seq data with DESeq2. Genome Biol. 15, 550 (2014).

100. H. Chen, VennDiagram: Generate High-Resolution Venn and Euler Plots. (2018), (available at https://cran.r-project.org/package=VennDiagram).

101. N. Gehlenborg, UpSetR: A More Scalable Alternative to Venn and Euler Diagrams for Visualizing Intersecting Sets. (2019).

102. F. A. Wolf, P. Angerer, F. J. Theis, SCANPY: Large-scale single-cell gene expression data analysis. Genome Biol. 19, 15 (2018).

103. C. Pardy, mpmi: Mixed-Pair Mutual Information Estimators. (2019), (available at https://cran.r-project.org/package=mpmi).

104. L. Scrucca, M. Fop, T. B. Murphy, A. E. Raftery, Mclust 5: Clustering, classification and density estimation using Gaussian finite mixture models. R J. 8, 289–317 (2016).

105. M. E. Futschik, L. Kumar, M. E Futschik, Introduction to Mfuzz package and its graphical user interface. Analysis, 1–13 (2009).

106. Z. Gu, L. Gu, R. Eils, M. Schlesner, B. Brors, Circlize implements and enhances circular visualization in R. Bioinformatics. 30, 2811–2812 (2014).

107. S. Lê, J. Josse, F. Husson, FactoMineR: An R package for multivariate analysis. J. Stat. Softw. 25, 1–18 (2008).

108. R. Suzuki, Y. Terada, H. Shimodaira, pvclust: Hierarchical Clustering with P-Values via Multiscale Bootstrap Resampling. (2019), (available at https://cran.r-project.org/package=pvclust).

109. R. M. Hope, Rmisc: Ryan Miscellaneous. (2013).

110. H. Hu, Y. R. Miao, L. H. Jia, Q. Y. Yu, Q. Zhang, A. Y. Guo, AnimalTFDB 3.0: A comprehensive resource for annotation and prediction of animal transcription factors. Nucleic Acids Res. 47, D33–D38 (2019).

111. A. Conesa, M. J. Nueda, A. Ferrer, M. Talón, maSigPro: A method to identify significantly differential expression profiles in time-course microarray experiments. Bioinformatics. 22, 1096–1102 (2006).

112. R Core Team, R: A Language and Environment for Statistical Computing (2020), (available at https://www.r-project.org/).

113. H. Wickham, M. Averick, J. Bryan, W. Chang, L. McGowan, R. François, G. Grolemund, A. Hayes, L. Henry, J. Hester, M. Kuhn, T. Pedersen, E. Miller, S. Bache, K. Müller, J. Ooms, D. Robinson, D. Seidel, V. Spinu, K. Takahashi, D. Vaughan, C. Wilke, K. Woo, H. Yutani, Welcome to the Tidyverse. J. Open Source Softw. 4, 1686 (2019).

114. M. Dowle, A. Srinivasan, data.table: Extension of ‘data.frame’ (2017).

115. D. Bates, M. Maechler, Matrix: Sparse and Dense Matrix Classes and Methods (2019).

116. M. Morgan, V. Obenchain, J. Hester, H. Pagès, SummarizedExperiment: SummarizedExperiment container (2019).

117. J. Baglama, L. Reichel, B. W. Lewis, irlba: Fast Truncated Singular Value Decomposition and Principal Components Analysis for Large Dense and Sparse Matrices (2019), (available at https://cran.r-project.org/package=irlba).

118. M. Maechler, P. Rousseeuw, A. Struyf, M. Hubert, K. Hornik, cluster: Cluster Analysis Basics and Extensions. (2019).

119. Z. Gu, R. Eils, M. Schlesner, Complex heatmaps reveal patterns and correlations in multidimensional genomic data. Bioinformatics. 32, 2847–2849 (2016).

120. R. Kolde, pheatmap: Pretty Heatmaps. (2019), (available at https://cran.r-project.org/package=pheatmap).

121. W. Chang, J. Cheng, J. Allaire, Y. Xie, J. McPherson, shiny: Web Application Framework for R (2020).

122. D. Attali, shinyjs: Easily Improve the User Experience of Your Shiny Apps in Seconds (2020), (available at https://cran.r-project.org/package=shinyjs).

123. F. Hahne, R. Ivanek, in Methods in Molecular Biology (Humana Press Inc., 2016; https://pubmed.ncbi.nlm.nih.gov/27008022/), vol. 1418, pp. 335–351.

124. N. Harmston, E. Ing-Simmons, M. Perry, A. Baresic, B. Lenhard, GenomicInteractions: R package for handling genomic interaction data (2020), (available at https://www.bioconductor.org/packages/release/bioc/html/GenomicInteractions.html).

